# Lipolysis of bone marrow adipocytes is required to fuel bone and the marrow niche during energy deficits

**DOI:** 10.1101/2022.03.21.485120

**Authors:** Ziru Li, Emily Bowers, Junxiong Zhu, Hui Yu, Julie Hardij, Devika P. Bagchi, Hiroyuki Mori, Kenneth T. Lewis, Katrina Granger, Rebecca L. Schill, Steven M. Romanelli, Simin Abrishami, Kurt D. Hankenson, Kanakadurga Singer, Clifford J. Rosen, Ormond A. MacDougald

## Abstract

To investigate roles for bone marrow adipocyte (BMAd) lipolysis in bone homeostasis, we created a BMAd-specific Cre mouse model in which we knocked out adipose triglyceride lipase (ATGL, *Pnpla2*). BMAd-*Pnpla2*^-/-^ mice have impaired BMAd lipolysis, and increased size and number of BMAds at baseline. Although energy from BMAd lipid stores is largely dispensable when mice are fed *ad libitum*, BMAd lipolysis is necessary to maintain myelopoiesis and bone mass under caloric restriction. BMAd-specific *Pnpla2* deficiency compounds the effects of caloric restriction on loss of trabecular bone, likely due to impaired osteoblast expression of collagen genes and reduced osteoid synthesis. RNA sequencing analysis of bone marrow adipose tissue reveals that caloric restriction induces dramatic elevations in extracellular matrix organization and skeletal development genes, and energy from BMAd is required for these adaptations. BMAd-derived energy supply is also required for bone regeneration upon injury, and maintenance of bone mass with cold exposure.

## Introduction

Bone marrow adipocytes (BMAds) are a heterogeneous cell population that form depots of bone marrow adipose tissue (BMAT) distinct from white, brown, and other adipose tissues (Z. Li, Hardij, Bagchi, Scheller, & MacDougald, 2018). Clinical associations generally demonstrate inverse relationships between BMAT and bone mass (Shen et al., 2007), or BMAT and circulating immune cells (Polineni et al., 2020), which may be due to the interactions between cells within the bone marrow niche. In addition to BMAds, the bone marrow niche contains osteoblasts, osteoclasts, hematopoietic cells, stromal/mesenchymal cells, blood vessels, and nerves (Vogler & Murphy, 1988). The relationships between BMAds, bone cells, and hematopoietic cells are influenced by their shared location within bone, an anatomically restricted system, such that expansion of one cell type is by necessity at the expense of others. For example, elevated BMAT is negatively correlated with low bone mass of aging and diabetes, whereas expansion of BMAT is associated with multiple hematopoietic disorders (Z. Li & MacDougald, 2021). Mechanistic links underlying these associations have proven challenging to investigate because they often involve complex intercellular, endocrine, and/or central mechanisms.

In addition to serving as an energy source, BMAds potentially influence the marrow niche through cell-to-cell contact, release of extracellular vesicles, and secretion of adipokines (e.g adiponectin) and cytokines (e.g. stem cell factor). Removal of these stimuli in mouse models of lipodystrophy, which lack BMAT, results in increased bone mass (Corsa et al., 2021; Zou et al., 2020; Zou et al., 2019). However, use of these models to investigate the direct effects of BMAT depletion is confounded by concurrent loss of white and brown adipose depots, which also regulate bone mass through myriad mechanisms, including secretion of adipokines (Riddle & Clemens, 2017; Zou et al., 2019). Although loss of BMAT in lipodystrophic mice was integral to the original finding that BMAT is a negative regulator of hematopoiesis, positive effects of BMAT on hematopoiesis have also been observed; these differences are attributed to use of distinct animal models and analysis of different skeletal sites (Ambrosi et al., 2017; Naveiras et al., 2009; Zhou et al., 2017). In rodents, there are two readily identifiable BMAd populations: constitutive BMAT (cBMAT) and regulated BMAT (rBMAT) (Scheller et al., 2015). cBMAT typically exists in distal tibia and caudal vertebrae, appears early in life, and has the histological appearance of white adipose tissue. rBMAT is found in proximal tibia and distal femur, and is comprised of single or clustered BMAds interspersed with hematopoietic cells. Whereas cBMAT generally resists change in response to altered physiological states, rBMAT expands with aging, obesity, diabetes, caloric restriction (CR), irradiation, and estrogen deficiency, and is reduced by cold exposure, fasting, β3-agonist, exercise, and vertical sleeve gastrectomy (Z. Li et al., 2018; Z. Li et al., 2019). These treatments also cause alterations to the skeleton and/or formation of blood cells, some of which may be secondary to effects on BMAds.

BMAds have long been believed to fuel maintenance of bone and hematopoietic cellularity because of their shared physical location in the marrow niche. However, this hypothesis has not been formally tested because current methods to target BMAds lack penetrance or cause recombination in other cell types, such as white adipocytes, osteoblasts, or bone marrow stromal cells, complicating the interpretation of interactions between BMAds and cells of the marrow niche. To circumvent this problem, we created a BMAd-specific Cre mouse model based on expression patterns of endogenous *Osterix* and *Adipoq*, and investigated roles for BMAds as a local energy source by deleting ATGL/*Pnpla2*, the rate-limiting enzyme of lipolysis. Consistent with *Pnpla2* deficiency in white and brown adipocytes (Ahmadian et al., 2011), BMAd-*Pnpla2*^-/-^ mice have impaired BMAd lipolysis, resulting in hypertrophy and hyperplasia of BMAT. Despite significant increases in bone marrow adiposity, hematopoietic abnormalities of BMAd-*Pnpla2*^-/-^ mice are negligible under basal conditions. However, the recovery of bone marrow myeloid lineages following sublethal irradiation is impaired with CR, and further reduced by BMAd-*Pnpla2* deficiency. Similarly, proliferative capacity of myeloid progenitors is also decreased with CR, and further inhibited in mice lacking BMAd-*Pnpla2*. Whereas alterations in bone parameters were not observed in *ad libitum* fed BMAd-*Pnpla2*^-/-^ mice, bone loss occurred under conditions of elevated energy needs such as bone regeneration or cold exposure, or reduced energy availability such as CR. Reduction of bone mass in CR BMAd-*Pnpla2*^-/-^ mice is likely due to impaired osteoblast functions such as expression of extracellular matrix genes and creation of osteoid. Gene profiling reveals that *Pnpla2* deletion largely blocks CR-induced genes within pathways of extracellular matrix organization and skeletal development, indicating that BMAd-derived energy contributes to skeletal homeostasis under conditions of negative energy balance.

## Results

### Generation of a BMAd-specific Cre mouse model (BMAd-Cre)

Based on previous studies showing that *Osterix* traces to osteoblasts and BMAds, but not to white adipocytes (Chen & Long, 2013; Mizoguchi et al., 2014), we used CRISPR/Cas9 to create *Osterix-FLPo* mice with an in-frame fusion of *Osterix* and optimized *FLPo*, separated by a *P2A* self-cleaving sequence to allow independent functioning of the two proteins (Figure 1A). To validate tissue-specific expression and FLPo efficiency, we bred *Osterix-FLPo* mice to FLP-dependent EGFP reporter mice, and observed EGFP-positive osteocytes, osteoblasts, BMAds, and a subset of marrow stromal cells within the bone (Figure 1B and 1C).

**Figure 1.**
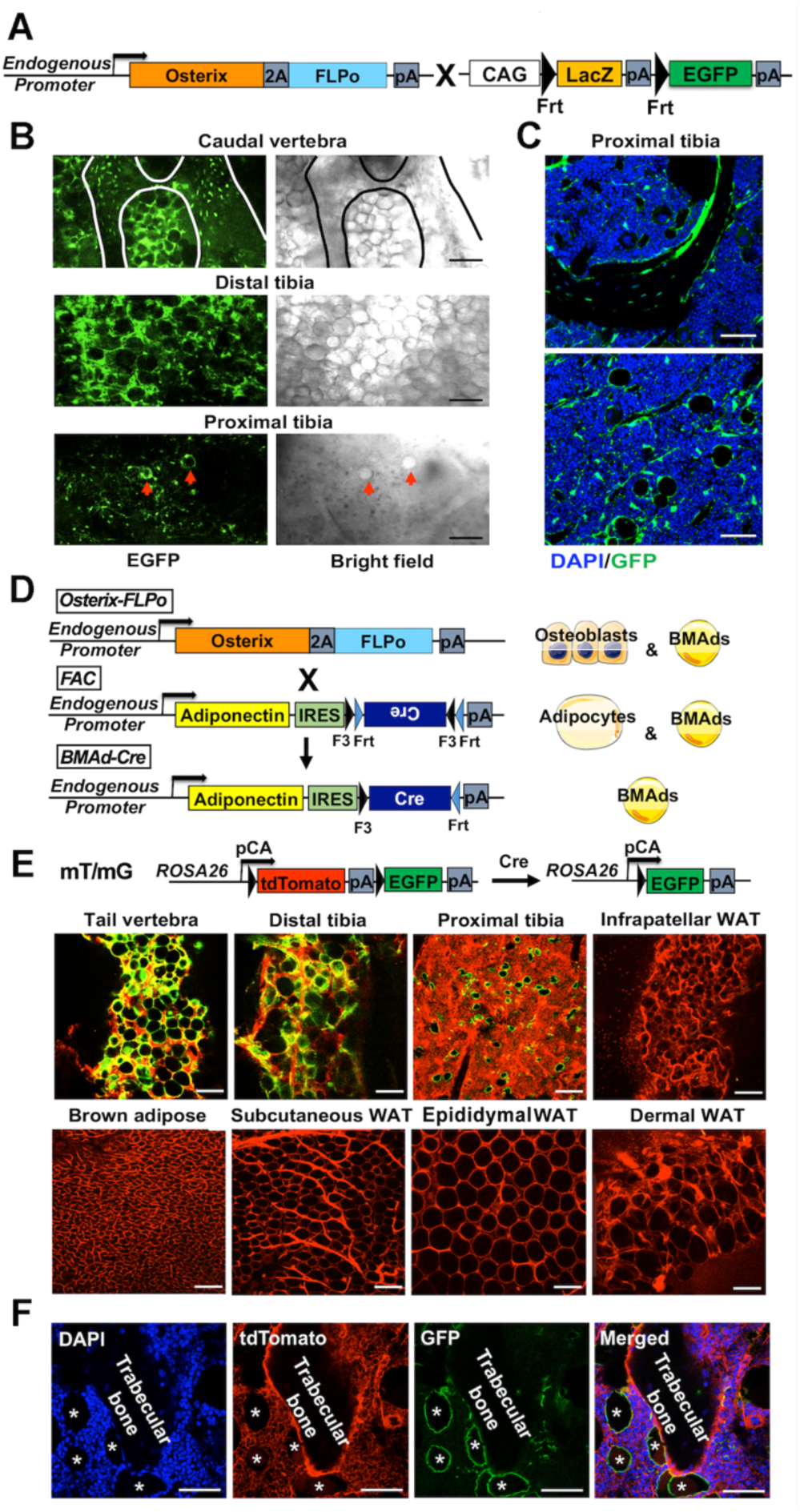
Generation of a BMAd-specific Cre mouse model (BMAd-Cre). A. Efficacy of *Osterix*-*FLPo* evaluated by crossing with FLP-dependent EGFP reporter to yield *Osterix*-EGFP. B. *Osterix-*EGFP male mice at 16 weeks were sacrificed. Fresh tissue confocal microscopy was performed on bisected caudal vertebrae, distal tibia and proximal tibia. Red arrows indicate singly dispersed BMAds. Scale bar; 100 μm. C. Frozen-sections of proximal tibiae slides from *Osterix*-EGFP were stained with Anti-GFP (green) and DAPI (blue). Scale bar; 50 μm. D. Schematic of how *Osterix*-*FLPo* recombines FLPo-activated *Adipoq-Cre* (FAC) in BMAd-Cre mice to restrict expression of Cre to BMAds. E. BMAd-Cre mice were bred with mT/mG reporter mice and resulting BMAd-mT/mG mice were sacrificed at 16 weeks of age and cellular fluorescence evaluated by fresh tissue confocal microscopy. Scale bar; 100 μm. F. Proximal tibia sections from BMAd-mT/mG mice were stained with antibodies to tdTomato, (red) and EGFP (green), and nuclei were counterstained with DAPI (blue). * indicates BMAds. Scale bar; 50 μm. <u>Figure 1 is associated with 2 supplements: Figure 1 - figure supplement 1; Figure 1 - figure supplement 2.</u>

We next created F<u>LPo</u>-dependent *<u>A</u>dipoq-<u>C</u>re* (FAC) mice, which contain an internal ribosome entry sequence (IRES) followed by FLPo-dependent *Cre* in reverse orientation within the 3’-untranslated region (UTR) of endogenous *Adipoq* gene (Figure 1D). FLPo expressed from the *Osterix* locus recombines *Cre* to the correct orientation in progenitors of osteoblasts and BMAds. However, because *Adipoq* is selectively expressed in adipocytes (Eguchi et al., 2011), Cre is expressed in BMAds, but not in osteoblasts or other adipose depots. Consistent with this schema, *Cre* in the correct orientation (flipped band, Figure 1 - figure supplement 1A-1C) is only observed in caudal vertebral DNA of mice positive for at least one copy each of *Osterix* (Mut band) and FAC (Ori band). The correct insertion of the sequence in *Adipoq* 3’-UTR was validated by Sanger sequencing following genomic PCRs that spanned endogenous *Adipoq* sequences, homology arms, IRES and Cre (Figure 1 - figure supplement 1D). To investigate the cell type-specificity of Cre activity, we next bred *BMAd-Cre* mice to mT/mG reporter mice (Muzumdar, Tasic, Miyamichi, Li, & Luo, 2007) (Figure 1E), in which all tissues and cells express red fluorescence (membrane-targeted tdTomato; mT) at baseline, and will express membrane-targeted EGFP in the presence of cell-specific Cre. We observed loss of tdTomato and gain of EGFP in BMAT depots of caudal vertebrae (tail) and tibiae, but not in brown or white adipose depots, or other tissues/organs such as liver, pancreas, muscle and spleen (Figure 1E and Figure 1 - figure supplement 1E). The correct orientation of *Cre* (flipped band) was also observed in mRNA isolated from distal tibiae and caudal vertebrae, but not WAT depots, of *Osterix-FLPo* mice positive for FAC (Figure 1 - figure supplement 1F). To evaluate conditions optimal for Cre-induced recombination, we visualized conversion of tdTomato to EGFP in *BMAd-Cre* mice at various ages and with different FAC copy numbers (Figure 1 - figure supplement 1G-1H) and found that the proportion of EGFP-positive BMAds increases with age and number of FAC alleles. Indeed, Cre efficiency is ∼80% in both male and female mice over 16 weeks of age with one Cre allele, and over 90% at 12 weeks of age in mice with two Cre alleles.

Previous studies found that a randomly inserted *Adipoq-Cre* bacterial artificial chromosome (BAC) causes recombination in osteoblasts (Bozec et al., 2013; Eguchi et al., 2011; Mukohira et al., 2019); thus, we tested whether this observation is true in *BMAd-Cre* mice, which rely on endogenous *Adipoq* expression that is restricted by *Osterix*. When *BMAd-Cre* mice are bred to mT/mG reporter mice, only tdTomato-positive cells are detectable on trabecular bone surfaces (Figure 1F), suggesting that osteoblasts are not targeted. However, labeling of a small subset of stromal/dendritic cells is consistently observed, likely due to expression of *Adipoq* within this cell population (Mukohira et al., 2019).

Although insertion of the FLPo-activated Cre cassette into the 3’-UTR of endogenous *Adipoq* overcomes potential problems that arise from random genomic integration, we considered the possibility that placement of the IRES-Cre cassette within the 3’-UTR might alter the expression and/or secretion of adiponectin. Although insertion of the IRES-Cre cassette may cause a slight reduction in mRNA expression in whole bone, the adiponectin mRNA in white adipose tissue is elevated by more than 3 fold (Figure 1 - figure supplement 2A). Thus, the cassette itself does not limit expression of mRNA. Instead, it appears that the 2 kb IRES-Cre cassette within the 3’-UTR impairs translatability of the mRNA since expression of adiponectin protein in BMAT and WAT is decreased by ∼50% (Figure 1 - figure supplement 2B and 2C). Circulating adiponectin concentrations appear to be decreased even further (Figure 1 - figure supplement 2D-2F), perhaps suggesting that flux of translated adiponectin protein is impaired through the secretion pathway. Of note, hypoadiponectinemia is associated with insulin resistance (Cook & Semple, 2010; N. Li et al., 2021), with discrepant reports on osteogenesis (Lewis, Edwards, Naylor, & McGettrick, 2021). Thus, we evaluated systemic metabolism in BMAd-Cre mice and found that body weight, glucose tolerance, WAT depot weights, and tibial trabecular and cortical bone variables are unaffected by this degree of hypoadiponectinemia (Figure 1 - figure supplement 2G-2P). To minimize variability between treatments, all mice used in the following studies were positive for both *Osterix-FLPo* and FAC. Gene knockout mice and their controls were determined by the presence or absence of floxed alleles, respectively.

### Ablation of adipose triglyceride lipase (ATGL, *Pnpla2*) causes BMAT expansion

A fundamental function of adipocytes is to store excess energy as triacylglycerols and to release non-esterified fatty acids and glycerol during times of negative energy balance. ATGL is the first and rate-limiting enzyme in the lipolytic process; thus, to determine the physiological functions of BMAd lipolysis in bone metabolism and hematopoiesis, we generated BMAd-*Pnpla2*^-/-^ mice, in which the gene encoding ATGL, *Pnpla2*, is specifically knocked out in BMAds. We validated the deletion of ATGL in caudal vertebrae, because cBMAT is abundant in this location. Mutant *Pnpla2* mRNA is observed in caudal vertebrae of BMAd-*Pnpla2* mice, but not in WAT (Figure 2A). Possible sources for the remaining wildtype (WT) *Pnpla2* signal include periosteal WAT, non-adipocyte cells, or perhaps from incomplete deletion of *Pnpla2* in BMAds. Immunofluorescent staining confirms loss of ATGL in proximal tibial rBMAds of BMAd-*Pnpla2*^-/-^ mice (Figure 2B). ATGL protein is reduced substantially in caudal vertebrae, but not in subcutaneous WAT (Figure 2C), which further confirms specificity of BMAd-Cre recombinase for BMAT.

**Figure 2.**
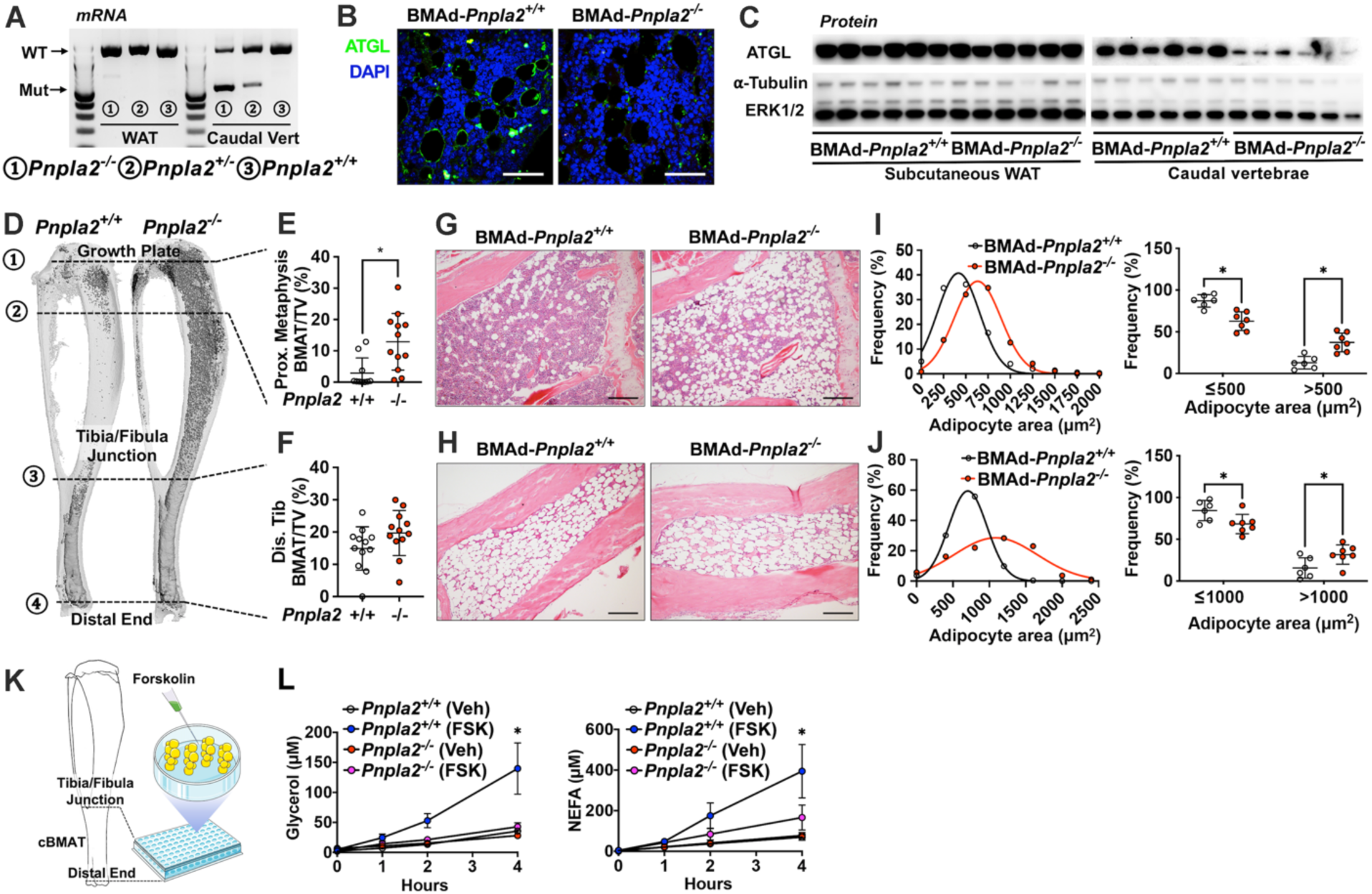
Ablation of adipose triglyceride lipase (ATGL; gene name *Pnpla2*) increases size and number of BMAd. A-J. Male mice of the indicated genotypes at 24 weeks of age were euthanized for investigation of white adipose tissue (WAT) and bone. A. RNA was extracted from WAT and caudal vertebrae and converted to cDNA. PCR products for wildtype (WT; 1257bp) *Pnpla2* and exon 2-7 knockout (Mut; 553 bp) bands were visualized. B. Decalcified proximal tibiae were sectioned and used for immunofluorescent staining for ATGL (green) expression. Slides were counterstained with DAPI (blue) for nuclei. Scale bar; 50 μm. C. Immunoblot analyses of ATGL, *α*-tubulin, and ERK1/2 in lysates from subcutaneous WAT and caudal vertebral. D-F. Decalcified tibiae were stained with osmium tetroxide and visualized by μCT (D). BMAT of proximal (E) and distal (F) tibia was quantified. Data are expressed as mean ± SD. * indicates *P* < 0.05 with a two-sample *t*-test. G-H. Decalcified tibiae were paraffin-sectioned and stained with Hematoxylin & Eosin. Representative pictures were taken from proximal (G) and distal (H) tibia. Scale bar; 200 μm. I-J. BMAd sizes from proximal (I) and distal (J) tibiae were quantified with MetaMorph software. Data are expressed as mean ± SD. * indicates *P* < 0.05 with two-way ANOVA with Sidak’s multiple comparisons test. K-L. Distal tibial BMAT was flushed from female BMAd-*Pnpla2*^-/-^ and their wildtype littermates at 24 weeks of age. For each n, distal tibial explants from 2 mice were combined per well and cultured in 2% BSA-HBSS solution (K). Subgroups from each genotype were treated with forskolin (FSK, 5 μM) or vehicle (Veh, DMSO). Released glycerol and non-esterified fatty acid (NEFA) in culture media at indicated time points were measured by colorimetric assay (n = 3-4 per treatment) (L). * indicates *Pnpla2*^+/+^ (FSK) different from *Pnpla2^+/+^* (Veh) and from *Pnpla2*^-/-^ (FSK) with *P* < 0.05 with two-way ANOVA with Sidak’s multiple comparisons test.

To quantify effects of *Pnpla2*-deficiency on bone marrow adiposity and cellularity, we used osmium tetroxide, and hematoxylin and eosin staining to evaluate BMAT quantity and cellular details, respectively. We found that BMAT volume is significantly increased in proximal tibiae and throughout the endocortical compartment of BMAd-*Pnpla2*^-/-^ mice, whereas differences in BMAT volume are not observed in distal tibiae, which is almost completely occupied by BMAT in WT mice (Figure 2D-2F). Histological analyses reveal increased BMAd number in proximal tibiae of BMAd-*Pnpla2*^-/-^ mice (Figure 2G and 2H). Depletion of *Pnpla2* also causes BMAd hypertrophy in both proximal and distal tibia, with an increased proportion of BMAds larger than 500 μm^2^ observed in proximal tibia, and larger than 1000 μm^2^ observed in distal tibia (Figure 2I and 2J). Secretion of basal glycerol and non-esterified fatty acids (NEFA) from *ex vivo* cultured explants of distal tibial BMAT was not different between genotypes. Whereas forskolin treatment greatly increased lipolysis in BMAT explants from control mice, secretion of glycerol and NEFA from BMAd-*Pnpla2*^-/-^ explants remained unchanged from basal rates (Figure 2K and 2L). These results confirm that BMAds of BMAd-*Pnpla2*^-/-^ mice have impaired lipolysis.

### BMAd lipolysis is required to maintain bone homeostasis in male mice under conditions of CR, but not when mice are fed *ad libitum*

We next evaluated whether loss of BMAd lipolysis in male BMAd-*Pnpla2*^-/-^ mice is sufficient to influence systemic physiology, or function of bone cells within the marrow niche. In mice fed normal chow *ad libitum*, we did not observe differences in body weight, glucose tolerance, or weights of soft tissue, including subcutaneous WAT (sWAT), epididymal WAT (eWAT) and liver (Figure 3 - figure supplement 1A-1E), which further confirms the tissue-specificity of our BMAd-knockout model. As observed above, *Pnpla2* deficiency causes expansion of proximal tibial rBMAT (Figure 3 - figure supplement 1F and 1G). Interestingly, when dietary energy is readily available, trabecular bone volume fraction, bone mineral density and trabecular number tend to be lower in BMAd-*Pnpla2*^-/-^ mice, but no significant differences were observed (Figure 3A-3C). These data also indicate that expansion of BMAT is not sufficient to cause bone loss, and that the correlation between elevated fracture risk and expansion of BMAT under a variety of clinical situations is not necessarily a causal relationship.

**Figure 3.**
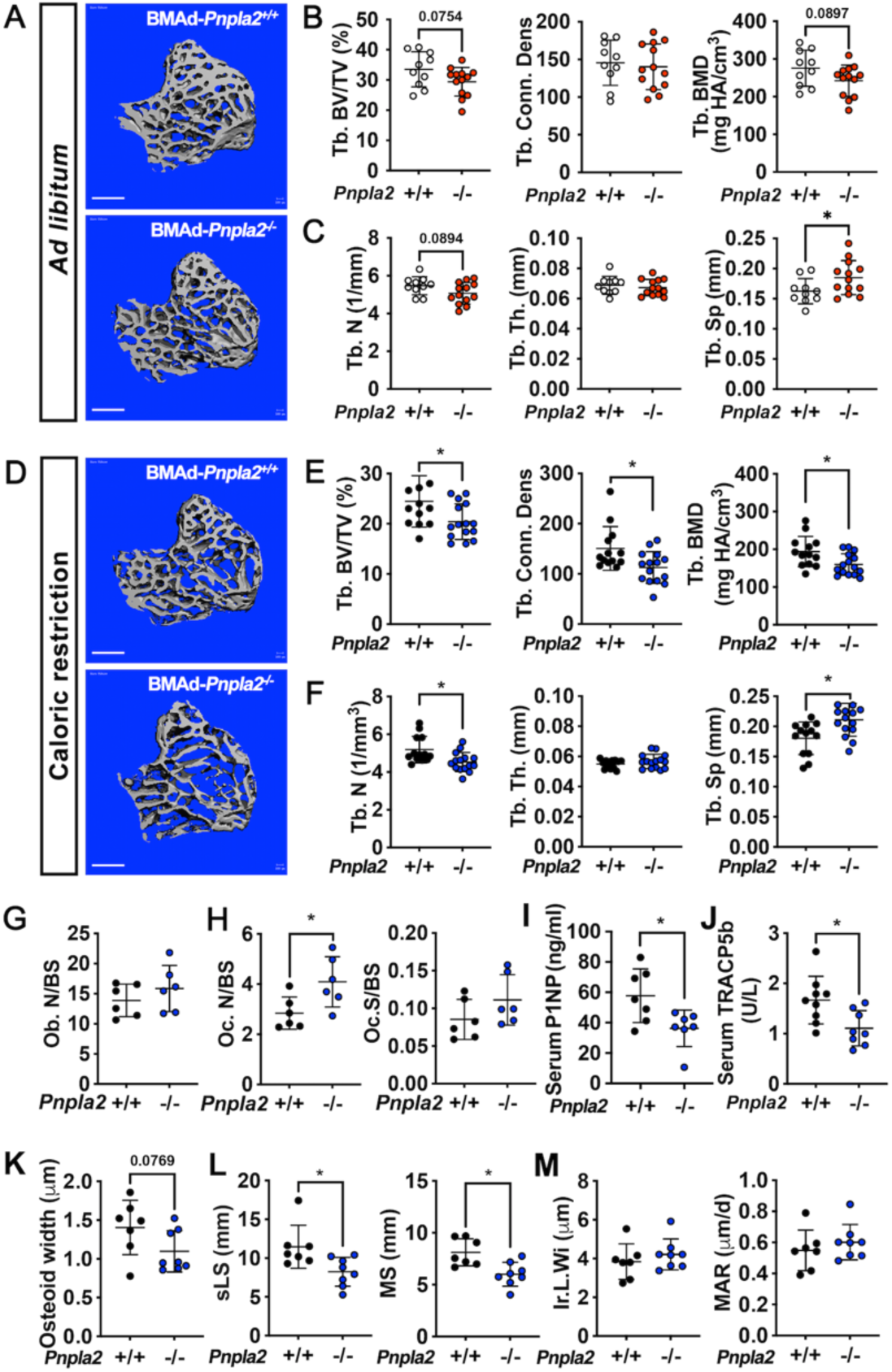
BMAd lipolysis is required to maintain bone homeostasis in male mice under CR conditions, but not when mice are fed *ad libitum*. A-C. Male BMAd-*Pnpla2*^-/-^ and their BMAd-*Pnpla2*^+/+^ littermates with *ad libitum* feeding were euthanized at 24 weeks of age. Tibiae from *ad libitum* mice were analyzed by μCT for indicated trabecular bone variables. Scale bars indicate 500 μm. D-M. Male mice at 18 weeks of age underwent 30% CR for 6 weeks. Two independent age- and sex-matched cohorts were plotted together for μCT parameters (D-F), one of those two cohorts was used for ELISA, and static or dynamic histomorphometry (G-M). D-F. Tibiae from CR mice were analyzed by μCT for indicated trabecular bone variables. Scale bar; 500 μm. G-H. Static histomorphometry analyses were performed to calculate osteoblast number (Ob. N), osteoclast number (OC. N) and osteoclast surface (Oc. S) per bone surface (BS). I-J. Concentrations of circulating P1NP and TRACP5b in CR mice. K. Osteoid quantification was performed on undecalcified plastic sections with Goldner’s Trichrome staining. L-M. Dynamic histomorphometry was performed on calcein-labelled trabecular bone from proximal tibia. sLS: single-labelled surface; MS: mineralized surface; Ir.L.Wi: inter-label width; MAR: mineral apposition rate. Data are expressed as mean ± SD. * indicates *P* < 0.05 with a two-sample *t*-test. <u>Figure 3 is associated with 4 supplements: Figure 3 - figure supplement 1; Figure 3 - figure supplement 2; Figure 3 - figure supplement 3; Figure 3 - figure supplement 4.</u>

Our experiments in *ad libitum* fed mice suggest that energy released from BMAds is dispensable for bone cell functioning when dietary energy is plentiful, which is supported by the comparable concentrations of glycerol and NEFA in circulation and in bone marrow supernatant in BMAd-*Pnpla2*^+/+^ and BMAd-*Pnpla2*^-/-^ mice (Figure 3 - figure supplement 1H and 1I). To determine whether male BMAd-*Pnpla2*^-/-^ mice have impaired bone homeostasis when dietary energy is limited, we next challenged mice with a 30% CR for six weeks. BMAd- *Pnpla2*^-/-^ mice did not have altered body weight, glucose tolerance, or tissue weights comparing with their controls (Figure 3 - figure supplement 1J-1N), suggesting that BMAds are not a critical source of circulating energy under these conditions. Although *ad libitum* fed BMAd-*Pnpla2*^-/-^ mice have increased rBMAT compared to controls (Figure 3 - figure supplement 1F and 1G), CR stimulates rBMAT expansion in control mice such that levels are comparable between genotypes (Figure 3 - figure supplement 1O and 1P). Interestingly, despite this similarity in BMAT volumes, BMAd-*Pnpla2*^-/-^ mice fed CR diet have reduced trabecular bone volume fraction, connective density, and bone mineral density when compared to controls (Figure 3E). This reduction in bone mass is due to decreased trabecular bone number and increased trabecular spacing, without effects on trabecular thickness (Figure 3D-3F). BMAds are a source of NEFA for vicinal osteoblasts (Maridas et al., 2019); thus, the reduction in trabecular bone in CR BMAd-*Pnpla2*^-/-^ mice likely results from impaired osteoblast function due to insufficient energy available in the bone marrow niche. Consistent with this notion, we observed a trend (P = 0.07) towards reduced NEFA concentrations in bone marrow supernatant of CR BMAd-*Pnpla2*^-/-^ mice, whilst circulating glycerol and NEFA were not different between genotypes (Figure 3 - figure supplement 1Q and 1R). Of note, changes in cortical bone area and thickness were not observed in BMAd- *Pnpla2*^-/-^ mice with either *ad libitum* or CR diet (Figure 3 - figure supplement 2A-2D), perhaps because cortical bone surfaces are less active.

To investigate mechanisms underlying trabecular bone loss in CR BMAd-*Pnpla2*^-/-^ mice, we next measured circulating markers of bone cell activity and performed histomorphometry. Osteoblast and osteoclast numbers are not changed by BMAd-*Pnpla2* deficiency under *ad libitum* conditions (Figure 3 - figure supplement 2E and 2F). Although circulating markers of bone formation (P1NP) and osteoclast activation (RANK Ligand) are not altered in BMAd- *Pnpla2*^-/-^ mice, markers for bone resorption (CTX-1) and osteoclast activation (TRACP5b) are decreased (Figure 3 - figure supplement 2G-2J). Although with CR, osteoblast numbers are not altered by a deficiency of BMAT lipolysis, the bone formation marker P1NP is decreased (Figure 3G and 3I), suggesting that osteoblast functions are impaired. Osteoclast numbers are increased in CR BMAd-*Pnpla2*^-/-^ mice, but the osteoclast surface per bone surface (Oc.S/BS) is not changed and osteoclast activation marker, TRACP5b, is decreased (Figure 3H and 3J), suggesting the osteoclast functions are inhibited. No differences were observed in RANK Ligand and CTX-1 with CR (Figure 3 - figure supplement 2K and 2L). We then visualized and quantified osteoid thickness with Goldner’s trichrome staining and observed a trend towards thinner osteoid layers without affecting osteoid surface (Figure 3K and Figure 3 - figure supplement 2M), which could be secondary to impaired secretion of collagen matrix by osteoblasts or to enhanced bone mineralization. Further, we injected mice with calcein to label those bone surfaces undergoing active mineralization. Dynamic histomorphometry data suggests that BMAd-*Pnpla2*^-/-^ mice have less bone-forming surface, as indicated by reduced single-labelled surface and mineral surface (Figure 3L). There were no differences in double-labelled bone surface, inter-label width, mineral apposition rate or osteoid maturation time (Figure 3M and Figure 3 - figure supplement 2N), suggesting that bone mineralization is not affected by loss of BMAd lipolysis. Taken together, these data support a model in which reduced trabecular bone in BMAd-*Pnpla2*^-/-^ mice is likely due to impaired ability of osteoblasts to secrete osteoid.

### BMAd lipolytic deficiency impairs myelopoiesis during regeneration

A single rBMAd connects to almost 100 hematopoietic cells (Robles et al., 2019), suggesting that BMAds potentially serve as important local energy sources for hematopoiesis. However, when fed *ad libitum*, male BMAd-*Pnpla2*^-/-^ mice do not exhibit differences in mature blood cells populations, as assessed by complete blood cell counts (Table 1) and bone marrow flow cytometry for hematopoietic populations (Table 2 and Figure 4 - figure supplement 1A and 1B). Although most hematopoietic stem and progenitor cells (HSPCs) do not depend on BMAd lipolysis, reduced numbers of granulocyte-monocyte progenitors (GMP) are observed in marrow of BMAd-*Pnpla2*^-/-^ mice fed *ad libitum* (Table 2). Of note, with CR, reduced numbers of circulating and bone marrow mature neutrophils are observed in BMAd-*Pnpla2*^-/-^ mice, although GMP numbers, which are the progenitor population, are not altered in CR BMAd-*Pnpla2*^-/-^ mice (Tables 1 and 2), perhaps due to impaired maturation of granulocytes. These data suggest that although myeloid cell defects are observed in BMAd-*Pnpla2*^-/-^ mice, other hematopoietic cell populations can metabolically compensate for the lack of BMAd lipolytic products.

To investigate further whether hematopoiesis is dependent upon BMAd lipolysis, we administered a sublethal dose of whole-body irradiation, and evaluated hematopoietic cell recovery. Following irradiation, white and red blood cell depletion and recovery were monitored by complete blood cell counts every 2-3 days (Figure 4 - figure supplement 1C-1E). In *ad libitum* fed mice, HSPCs (Figure 4 - figure supplement 1F and 1G) and mature/immature hematopoietic cells (Figure 4A-4H) are not influenced by deficiency of BMAd-lipolysis. However, CR alone decreases bone marrow cellularity (Figure 4A), and HSPC (Figure 4 - figure supplement 1F and 1G) and neutrophil numbers, without affecting monocytes and lymphocytes (Figure 4B-4E). In CR BMAd-*Pnpla2^-/-^* mice, HSPCs are not altered compared to CR controls, but bone marrow cellularity, monocytes and neutrophils are decreased further. Preneutrophils and immature neutrophils are also reduced by CR, and show additional decline in CR mice lacking BMAd-*Pnpla2* (Figure 4F-4G), but mature neutrophils are only affected by CR (Figure 4H). These reductions in monocytes and neutrophils suggest that BMAd-lipolysis is required for myeloid cell lineage regeneration when energy supply from circulation is limited. CFU assays, which are optimized for the growth of myeloid progenitor cells (CFU-Granulocytes, CFU-Macrophages and CFU-GM), demonstrate fewer granulocyte progenitor colonies (CFU-G) from CR BMAd-*Pnpla2^-/-^* mice (Figure 4I). The proliferative capacity of macrophage progenitors (CFU-M) is impaired when derived from BMAd-*Pnpla2*^-/-^ mice fed *ad libitum*, or when derived from CR mice of either genotype (Figure 4J). CR marrow produced fewer granulocyte-macrophage progenitors (CFU-GM), which was further reduced when isolated from mice with impaired BMAd-lipolysis (Figure 4K). These data suggest that proliferative and differentiation capacities of cultured myeloid cell progenitors may have been reprogrammed in the marrow niche when their energy supply is restricted either by diet or impaired BMAd lipolysis (Figure 4L).

**Figure 4.**
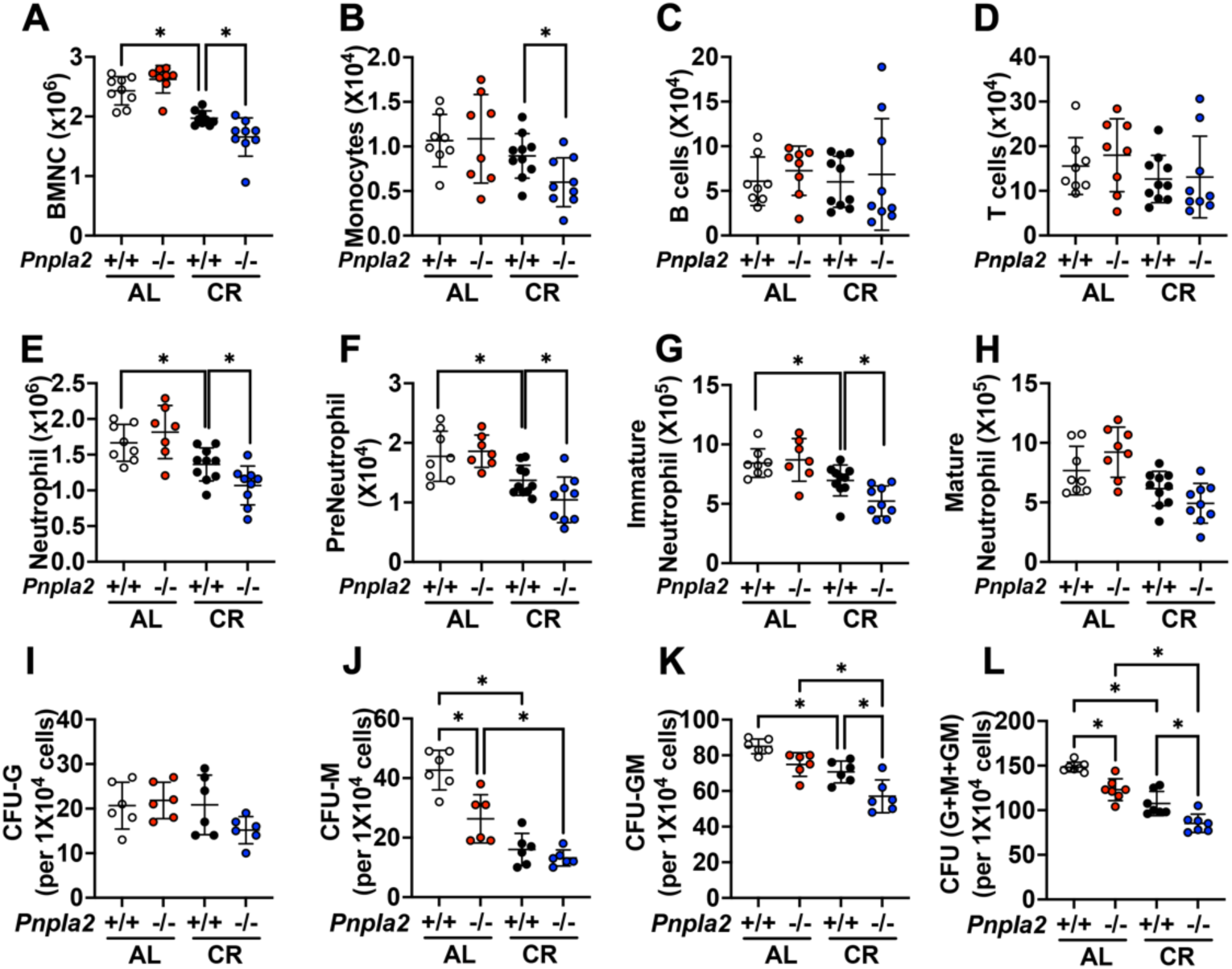
BMAd-*Pnpla2* deficiency impairs myelopoiesis. A-H. BMAd-*Pnpla2*^-/-^ mice and littermate controls (*Pnpla2*^+/+^) were caloric restricted (CR) for 20 weeks or remained on an *ad libitum* (AL) diet, and then received whole-body irradiation (6 Gy). Mice were euthanized 9 days post-irradiation. Femurs were collected for flow cytometry to measure the regeneration of hematopoietic cells. Bone marrow mononuclear cells (BMNCs), monocytes, B and T lymphocytes and neutrophils were quantified. I-L. CFU assays. Femora and tibial bone marrow cells were isolated from BMAd-*Pnpla2*^-/-^ mice and littermate controls (*Pnpla2*^+/+^), which had been fed ad libitum (AL) or a caloric restricted (CR) diet for 20 weeks. After counting, 1x10^4^ cells were plated for CFU assays. Colonies were counted by an independent expert in a blinded manner 7 days after plating. Data are expressed as mean ± SD. * indicates *P* < 0.05 with one-way ANOVA with Tukey’s multiple comparisons test. <u>Figure 4 is associated with 1 supplement: Figure 4 - figure supplement 1.</u>

### BMAd lipolysis is not required to maintain bone homeostasis under calorie-restricted conditions in female mice

To determine whether sex influences responses of BMAd-*Pnpla2*^-/-^ mice to CR, we performed similar experiments in female mice at 20 weeks of age. As expected, after six weeks of CR, both control and knockout mice exhibited comparable reduced body weights, random blood glucose concentrations, and tissue weights (Figure 3 - figure supplement 3A and 3B). Although CR and *Pnpla2* deficiency cause expansion of proximal tibial rBMAT (Figure 3 - figure supplement 3C), neither CR nor BMAd *Pnpla2* deletion cause alterations in trabecular or cortical bone variables in female mice (Figure 3 - figure supplement 3D and 3E). Consistent with these findings, previous studies showed that CR adult or aged female mice have milder bone loss than in males (Z. Li & MacDougald, 2021). Although male BMAd-*Pnpla2*^-/-^ mice exhibited deficiencies in circulating neutrophils, experiments with female mice did not reveal differences in white or red blood cell populations (Figure 3 - figure supplement 3F). We next considered whether estrogen protects female mice from CR-induced osteoporosis. Thus, we ovariectomized control and BMAd-*Pnpla2*^-/-^ mice two weeks prior to initiation of CR. Following initiation of CR, both control and knockout mice demonstrated rapid reduction in body weight for two weeks, then gradually stabilized during the following ten weeks (Figure 3 - figure supplement 4A). We did not observe significant differences in glucose tolerance or bone length with CR or *Pnpla2* deletion (Figure 3 - figure supplement 4B and 4C). Additionally, although CR caused reduction in tissue weights, these variables were not different between genotypes (Figure 3 - figure supplement 4D). Interestingly, despite the dramatic increases of BAMT responding to CR or BMAd- *Pnpla2* deficiency, bone volume fraction and trabecular number were increased by CR in ovariectomized mice (Figure 3 - figure supplement 4E and 4F), suggesting that the metabolic benefits of CR diet combat detrimental effects of estrogen deficiency and/or aging. Although we observed a trend toward reduced trabecular thickness in *Pnpla2*-deficient mice, other skeletal parameters were not affected (Figure 3 - figure supplement 4F).

### Coupling of BMAd *Pnpla2* deletion and CR results in extensive alterations to the bone marrow transcriptome

To determine mechanisms by which BMAd *Pnpla2* deficiency causes bone loss in CR male mice, we profiled overall gene expression using bulk RNAseq in bone marrow plugs from distal tibiae, a skeletal location highly enriched with BMAT. PCA plots show that RNA profiles from CR groups are distinct from *ad libitum* controls (Figure 5 - figure supplement 1A). Whereas *Pnpla2* deficiency does not cause gene expression to diverge substantially in mice fed *ad libitum*, loss of *Pnpla2* interacts with CR to cause a well-segregated pattern of gene expression (Figure 5 - figure supplement 1A). By our criteria (padj < 0.05, |log2 fold change|>1), CR changes expression of 1,027 genes compared to *ad libitum* controls. Although *Pnpla2* deficiency alone only alters 10 genes in BMAd-*Pnpla2*^-/-^ mice fed *ad libitum*, loss of *Pnpla2* in CR mice causes alterations in 1,060 genes (Figure 5 - figure supplement 1B). Analyses of genes regulated in BMAd-*Pnpla2*^+/+^ mice with CR reveals four distinct clusters (Figure 5A). Approximately 80% of genes fall in cluster 1, which are upregulated by CR in control mice, with induction largely blocked by *Pnpla2* deficiency in CR mice. Pathway analyses of cluster 1 with Metascape (https://metascape.org) reveals that regulated genes are associated with vasculature development and cellular response to growth factor stimulus, which likely reflects adaptation mechanisms to compensate for energy insufficiency (Figure 5 - figure supplement 1C). In addition, genes associated with skeletal system development, extracellular matrix organization, and adipogenesis pathways are upregulated by CR in control mice, but these effects are blunted by *Pnpla2* deletion (Figure 5 - figure supplement 1C). Cluster 2 includes genes that are mildly upregulated by CR in control mice and are further increased with *Pnpla2*-deficiency. However, no specific pathways are enriched in this gene set (Figure 5 - figure supplement 1D). Cluster 3 highlights genes that are downregulated by CR treatment regardless of genotype (Figure 5 - figure supplement 1E), and which are associated with B cell proliferation and interleukin-8 production, which may partially explain the changes observed in hematopoietic cellularity. Cluster 4 contains 80 genes that are down-regulated independently by CR and *Pnpla2-* deficiency, and are associated with regulation of ossification and calcium-mediated signaling (Figure 5 - figure supplement 1F).

**Figure 5.**
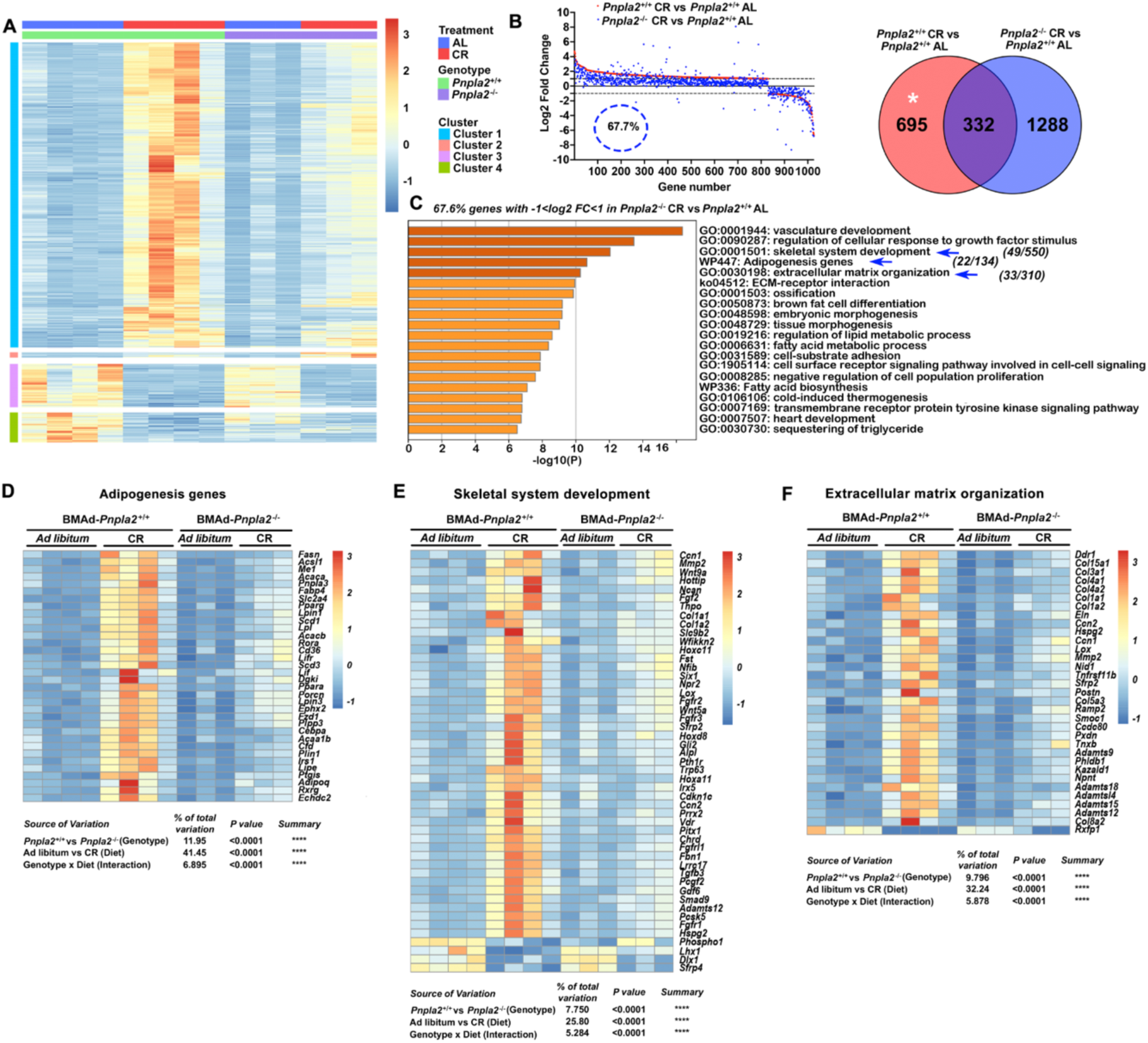
BMAd-*Pnpla2* deficiency causes extensive alterations to the bone marrow transcriptome only when coupled with CR. Male control and BMAd-*Pnpla2*^-/-^ mice at 24 weeks of age were either fed AL or underwent 30% CR for 6 weeks. Distal tibial cBMAT was flushed and cBMAT from two mice was pooled as one sample for RNAseq analyses (n of 3 or 4 per treatment). A. Differential genes with our criteria (P adj<0.05 and |Log2 fold change|>1) between BMAd-*Pnpla2*^+/+^ CR and BMAd-*Pnpla2*^+/+^ *ad libitum* (AL) were grouped into 4 clusters. B. Genes different between BMAd-*Pnpla2*^+/+^ CR and BMAd-*Pnpla2*^+/+^ AL were ordered from maximum to minimum log2 fold change (red dots), and compared to corresponding data from BMAd-*Pnpla2*^-/-^ CR versus BMAd-*Pnpla2*^+/+^ AL (blue dots). Venn diagram shows the differential genes between BMAd- *Pnpla2*^+/+^ CR versus BMAd-*Pnpla2*^+/+^ AL BMAT; and BMAd-*Pnpla2*^-/-^ CR versus BMAd-*Pnpla2*^+/+^ AL BMAT. C. Pathway analyses of genes significantly changed by CR in BMAd-*Pnpla2*^+/+^ mice, but not in CR mice lacking *Pnpla2* (indicated by * area in panel B). Pathways further analyzed by heatmap indicated with blue arrows. D-F. Expression Z-scores of genes related to adipogenesis (D), skeletal system development (E) and extracellular matrix organization (F) were shown as heatmap. Effects of genotype and diet, and their interactions were analyzed by three-way ANOVA. <u>Figure 5 is associated with 2 supplements: Figure 5 - figure supplement 1; Figure 5 - figure supplement 2.</u>

To evaluate further whether BMAd-*Pnpla2* is required for the adaptation of BMAT to CR, we graphically ordered genes from those maximally induced to those most repressed by CR in control mice (Figure 5B; red dots). Of these genes, 67.7% do not meet our criteria for regulated expression by CR in BMAd-*Pnpla2*^-/-^ mice (blue dots). Pathway analyses on the 32.3% of genes regulated by CR regardless of *Pnpla2* deficiency reveals association with ribonuclease activity, response to hormones and other transmembrane signaling pathways (Figure 5 - figure supplement 2A). Pathway analyses of genes for which *Pnpla2* is required for response to CR identifies vasculature development and cellular response to growth factor stimulus, similar to cluster 1, followed closely by skeletal system development, adipogenesis genes, and extracellular matrix organization pathways (Figure 5C). A heatmap shows that the adipogenic genes, including *Adipoq, Fabp4, Cebpa, Lipe, Lpl, Pparα, Pparγ, Scd1, Plin1, Cd36, Fasn* and *Acaca*, are upregulated by CR in control mice (Figure 5D), a subset of which were confirmed by qPCR (Figure 5 - figure supplement 2B). These changes may help explain molecular mechanisms underlying BMAT expansion following CR. It is important to note that whereas CR BMAd-*Pnpla2*^+/+^ and BMAd-*Pnpla2*^-/-^ mice had comparable amounts of BMAT (Figure 3 - figure supplement 1O and 1P), adipocyte gene expression is greatly suppressed in mice with BMAT lacking *Pnpla2 (Figure 5D)*. This observation is consistent with prior work showing that adipocyte-specific deletion of *Pnpla2* results in decreased expression of genes associated with lipid uptake, synthesis, and adipogenesis (Schoiswohl et al., 2015), perhaps because NEFA and associated metabolites act as PPAR ligands are decreased (Mottillo, Bloch, Leff, & Granneman, 2012). In addition, genes related to endogenous fatty acid biosynthesis are also increased by CR in BMAd-*Pnpla2*^+/+^ mice, but not in mice lacking *Pnpla2* (Figure 5 - figure supplement 2C). Interestingly, myeloid leukocyte differentiation genes follow a similar pattern (Figure 5 - figure supplement 2D), which may contribute to reduced neutrophil production and impaired myeloid cell proliferation of CR BMAd-*Pnpla2*^-/-^ mice (Tables 1 and 2, and Figure 4).

We previously observed increased bone loss in BMAd-*Pnpla2*^-/-^ mice challenged with CR (Figure 3). To investigate potential mechanisms underlying this bone loss, we analyzed pathways from Cluster 1 related to bone metabolism, skeletal system development and extracellular matrix organization. Interestingly, osteoblast-derived alkaline phosphatase (*Alpl*)*, Col1a1* and *Col1a2*, and bone marrow Fgf/Fgfr and Wnt signaling-related molecules are highly induced by CR in control mice but not in *Pnpla2*-deficient mice (Figure 5E); many of these genes are also found in the ossification pathway (data not shown). Multiple collagen genes, *Lox*, and *Adamts* (A Disintegrin and Metalloproteinase with Thrombospondin motifs) family members, which are multidomain extracellular protease enzymes and play key roles in extracellular matrix remodeling, are also upregulated by CR in control mice, with effects largely eliminated by *Pnpla2* deficiency (Figure 5F). These findings may partially explain why osteoid thickness tends to be thinner in BMAd-*Pnpla2^-/-^* mice (Figure 3K). In this regard, 20 collagen genes are significantly up-regulated in control mice following CR (Figure 5 - figure supplement 2E), whereas only four collagen genes are elevated in CR BMAd-*Pnpla2*^-/-^ mice. Taken together, these data suggest that under conditions of limited dietary energy, BMAds provide energy to maintain osteoblast functions, including the secretion of collagen matrix for osteoid synthesis.

### BMAd lipolysis is required for trabecular and cortical bone regeneration

To test whether BMAds are a critical source of local energy under conditions where energy needs are elevated, we investigated the impact of CR and BMAd *Pnpla2* deficiency on bone regeneration. To do this, we created a 0.7 mm hole in the proximal tibia, approximately 1 to 2 mm distal from the growth plate, and evaluated trabecular and cortical parameters nine days later (Figure 6A). As expected, CR impairs formation of new trabecular bone in the region of interest (ROI) by decreasing trabecular bone volume fraction, bone mineral density, and bone mineral content (Figure 6B). Importantly, these effects of CR are mimicked by BMAd-specific *Pnpla2* deletion; however, gene deletion does not compound effects of CR on impaired bone regeneration, perhaps because bone volume is already low with either CR or *Pnpla2* deficiency alone. We then visualized newly formed cortical bone with Safranin O/ Fast Green (SO/FG) staining of paraffin-embedded proximal tibia sections (Figure 6C). Of note, both bone marrow stromal and periosteal cells contribute to the cortical bone regeneration and are derived from common mesenchymal progenitors (Duchamp de Lageneste et al., 2018). Although deficiency of BMAd-lipolysis is unlikely to affect the periosteal cell functions when the cortical bone is intact, it may interact with this cell population during bone regeneration. CR also impaired formation of new cortical bone by decreasing cortical bone volume fraction, bone mineral density, and bone mineral content. As with trabecular bone formation, effects of CR on cortical bone are mimicked by BMAd-specific depletion of *Pnpla2*, but effects of gene deletion do not exacerbate CR-induced impaired bone regeneration (Figure 6D). Taken together, these data provide compelling evidence that under conditions where energy requirements are high, BMAd provide a critical local energy source for bone regeneration.

**Figure 6.**
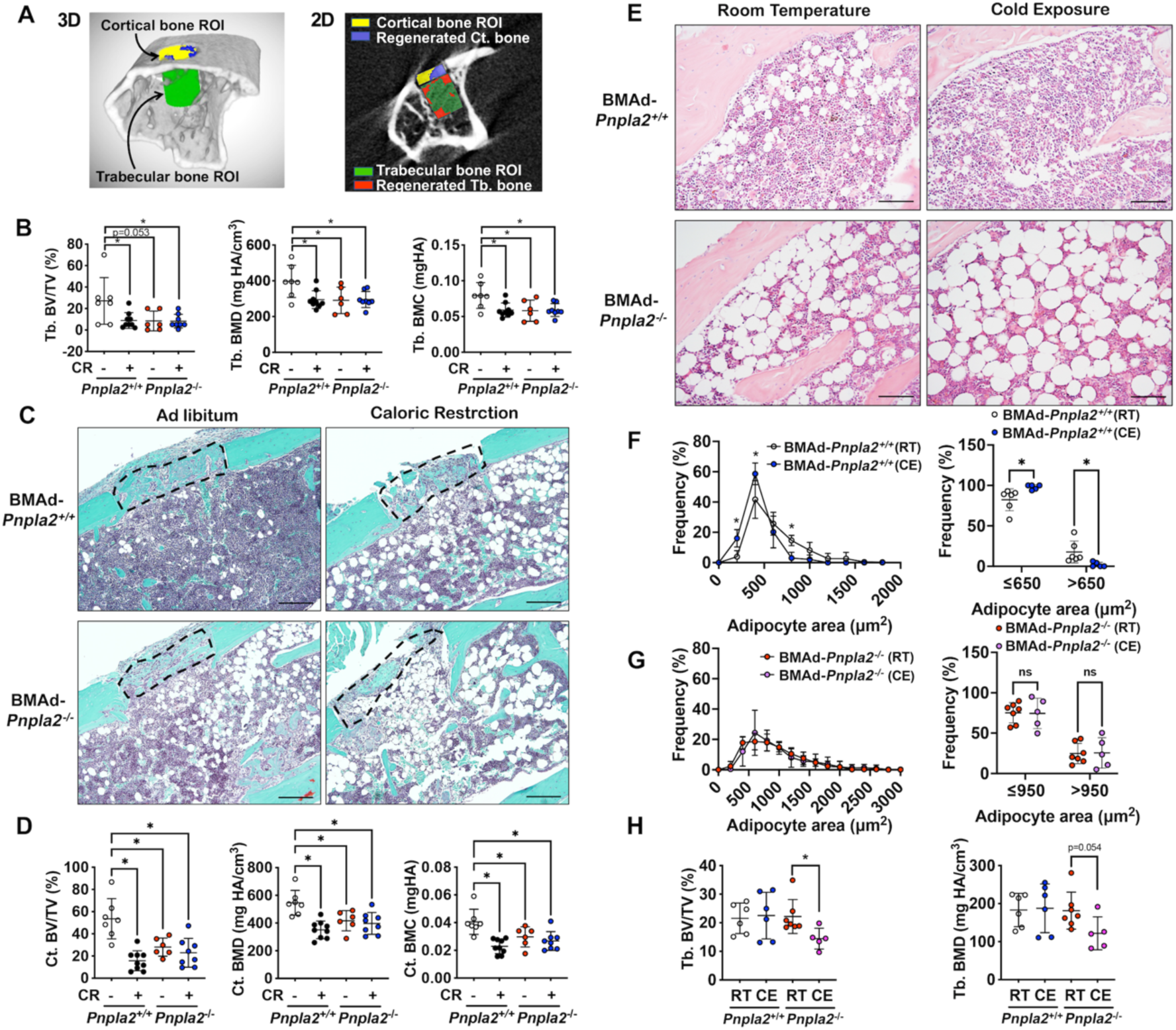
Energy from BMAd is required for trabecular bone regeneration and protects against bone loss caused by chronic cold exposure. A-D. BMAd-*Pnpla2*^+/+^ and BMAd-*Pnpla2*^-/-^ male mice at 24 weeks of age fed with chow diet (-) or underwent 30% CR for 6 weeks (+). A 0.7 mm proximal tibial defect was created 1 to 2 mm distal to the growth plate. Tibiae were collected 9 days after surgery. MicroCT was performed to analyze trabecular and cortical bone regeneration. A. Representative analyzing images of bone defect. Example of μCT showing trabecular bone region of interest (ROI; green) and newly generated trabecular bone (red); cortical bone ROI (yellow) and new- formed cortical bone (purple). B. Quantification of regenerated trabecular bone volume fraction (BV/TV), mineral density (BMD) and mineral content (BMC). Data are expressed as mean ± SD. * indicates *P* < 0.05 with one-way ANOVA with Tukey’s multiple comparisons test. C. Safranin O/ Fast Green (SO/FG) staining of new cortical bone formation in defect sites. Scale bar; 200μm. D. Quantification of regenerated cortical bone volume fraction (BV/TV), mineral density (BMD) and mineral content (BMC). Data are expressed as mean ± SD. * indicates *P* < 0.05 with one-way ANOVA with Tukey’s multiple comparisons test. E-I. Female BMAd-*Pnpla2*^+/+^ and BMAd-*Pnpla2*^-/-^ mice at 20 weeks of age were singly housed at 5°C for 3 weeks without enrichments. Tibiae were collected for sectioning and μCT analyses. E. Proximal tibiae were decalcified and paraffin-sectioned for H&E staining. Representative images for proximal tibia are shown. Scale bar, 100 μm. F-G. Quantification of BMAds from H&E-stained slides using MetaMorph software. Comparison of BMAd size at room temperature (RT) versus cold exposure (CE) in mice of indicated genotypes. Data are expressed as mean ± SD. * indicates *P* < 0.05 with two-way ANOVA with Sidak’s multiple comparisons test. H. Trabecular bone volume fraction (BV/TV) and mineral density (BMD) were quantified by μCT. Data are expressed as mean ± SD. * indicates *P* < 0.05 with one-way ANOVA with Tukey’s multiple comparisons test.

### Energy from BMAd protects against bone loss caused by chronic cold exposure

To explore further under what conditions BMAd lipolysis may be critical for bone cell functions, control and BMAd-*Pnpla2*^-/-^ female mice at 20 weeks of age were housed at room (22°C) or cold (5°C) temperatures for three weeks. Cold exposure is well-documented to increase energy expenditure and adaptive thermogenesis, largely fueled by energy stored in WAT depots. As expected (Scheller et al., 2015; Scheller et al., 2019), cold exposure results in smaller regulated BMAds within the proximal tibia of BMAd-*Pnpla2^+/+^* mice (Figure 6E and 6F). In contrast, BMAd size is increased at baseline in BMAd-*Pnpla2*^-/-^ mice, with no reduction observed with cold exposure (Figure 6E and 6G). These data indicate that intact BMAd lipolysis is required for reduction of BMAd size with cold exposure. We then evaluated bone mass by μCT and found that whereas trabecular bone of the proximal tibia is maintained in BMAd-*Pnpla2*^+/+^ mice with cold stress, trabecular bone volume fraction and bone mineral density decline with cold exposure in BMAd-*Pnpla2*^-/-^ mice (Figure 6I). These results indicate that lipolysis from vicinal BMAds is required for maintenance of bone when energy needs are high or when energy supply is limited.

## Discussion

To date, investigation of the physiological functions of BMAT have been hampered by the lack of a BMAd-specific mouse model. Although a number of BMAT depletion models show high bone mass, including A-ZIP (Naveiras et al., 2009), *Adipoq*-driven DTA (Zou et al., 2019), and *Adipoq*-driven loss of *Pparγ* (Wang, Mullican, DiSpirito, Peed, & Lazar, 2013), *Lmna* (Corsa et al., 2021), or *Bscl2* (McIlroy et al., 2018), these lipodystrophic mice also lack white and brown adipose depots and thus exhibit global metabolic dysfunction, including fatty liver, hyperlipidemia and insulin resistance. Of note, *Adipoq*-driven Cre used in those studies also causes recombination in bone marrow stromal cells (Mukohira et al., 2019). Similarly, mouse models or treatments that result in BMAT expansion, such as *Prx1*-driven *Pth1r* knockout mice (Fan et al., 2017), CR, thiazolidinedione administration, estrogen-deficiency, and irradiation (Z. Li & MacDougald, 2021), also cause bone loss, but again, effects on bone may be secondary to lack of promoter specificity or systemic effects. Lineage tracing studies have previously been performed to determine cell-specific markers for BMAds. For example, both *Prx1* and *Osterix* are restricted to bone and trace to 100% of BMAds, but are also expressed in mesenchymal cells (Logan et al., 2002; Mizoguchi et al., 2014). *Nestin* and leptin receptor (*LepR*) label over 90% of BMAds, but also trace to stromal cells (Zhou et al., 2017). Whereas Pdgf-receptor α (*PdgfRα)*-driven Cre expression causes recombination in all white adipocytes, only 50-70% of BMAds are traced (Horowitz et al., 2017). Thus, our strategy for targeting BMAds using dual expression of *Osterix* and *Adipoq* will be critical for improving interpretability of experiments on roles for BMAds in the marrow niche.

After generating this novel BMAd-specific Cre mouse model, we successfully ablated *Pnpla2* in BMAds, and demonstrated the necessity of ATGL in BMAd lipolysis. Our studies directly demonstrate for the first time the importance of BMAd lipolysis in myelopoiesis and bone homeostasis under conditions of energetic stress, including CR, irradiation, bone regeneration and cold exposure. With the loss of peripheral WAT in CR mice, there is a dramatic increase in BMAT throughout the tibia, and this is accompanied by increased expression of adipocyte genes, including those involved in lipid uptake, de novo lipogenesis, and lipolysis. BMAT expansion with CR has been observed in both rodents and humans (Cawthorn et al., 2014; Devlin et al., 2010; Fazeli et al., 2021), and mechanisms underlying this observation are still not fully understood. We speculate that CR promotes lipolysis in peripheral adipose tissues, and the released non-esterified fatty acids have increased flux to bone marrow, where they are used either directly to maintain hematopoiesis and bone homeostasis, or used indirectly after having been stored and released from BMAT. Thus, we speculate that BMAds have elevated rates of lipid uptake, lipogenesis and lipolysis with CR. However, when BMAd lipolysis is impaired, this dynamic cycle is halted, which is reflected by the blunted induction of adipocyte genes in response to CR. This BMAd quiescence causes a shortfall in local energy supply and contributes to hematopoietic cell and osteoblast dysfunction. In this regard, two major collagens secreted by osteoblasts, *Col1a1* and *Col1a2*, and *Alpl* were increased by CR in WT mice, but not in calorie-restricted BMAd-*Pnpla2^-/-^* mice. These data are consistent with the trend for thinner osteoid observed in CR BMAd-*Pnpla2*^-/-^ mice and reduced circulating concentrations of the bone formation marker P1NP. Although bone mineralization is not impaired, new bone forming surface is reduced in CR mice that lack BMAd lipolysis. In addition, the number of osteoclasts on trabecular bone surface is higher in CR BMAd-*Pnpla2*^-/-^ mice; however, osteoclast surface per bone surface and a circulating marker of bone turnover (CTX-1) are not increased. A serum marker for osteoclast activity, TRACP5b, is decreased in BMAd-*Pnpla2*^-/-^ mice either fed *ad libitum* or a CR diet. Taken together, our studies suggest that bone mass reduction in CR BMAd-*Pnpla2*^-/-^ mice is less likely due to degradation of bone by osteoclasts, and more likely due to impaired osteoblast function, including osteoid production.

Interestingly, the contributions of BMAd lipolysis to bone homeostasis appear to be more important in male mice compared to females. Although we considered that a stronger phenotype might be revealed in female mice following estrogen depletion, the low bone mass observed with ovariectomy or CR may represent a critical threshold that is strongly defended through mechanisms independent of BMAd lipolysis. Alternatively, it is possible that androgens increase energy requirements of bone such that male mice are more dependent on BMAd lipolysis under stressful conditions. Importantly, we observed that bone volume fraction and trabecular number were increased by 12 weeks of CR in OVX mice of both genotypes, suggesting that the metabolic and anti-aging benefits of CR somehow block the bone loss associated with estrogen deficiency as mice age to 40 weeks.

Surprisingly, despite expansion of BMAT in *ad libitum*-fed BMAd-*Pnpla2^-/-^* male and female mice, we did not observe differences in skeletal parameters. Indeed, the critical role of BMAd lipolysis in fueling osteoblast was observed in BMAd-*Pnpla2^-/-^* male mice only when energy needs were high, or the availability of dietary energy was low. Of note, cellular protein synthesis typically uses 25 to 30% of the oxygen consumption coupled to ATP synthesis (Rolfe & Brown, 1997), and this percentage may be higher in osteoblasts since protein synthesis and translation are integral to cell function. Additionally, fatty acids contribute substantially to the energy demands of bone tissue and cells (Adamek, Felix, Guenther, & Fleisch, 1987), and in the absence of BMAd lipolysis, these energy needs must be met from circulating lipids, glucose, lactate and amino acids. Uptake of fatty acids by osteoblasts is likely mediated by the CD36 fatty acid translocase, which is required in mice to maintain osteoblast numbers and activity (Kevorkova et al., 2013). Further, impairment of fatty acid oxidation in osteoblasts and osteocytes led to reduced postnatal bone acquisition in female, but not male mice. Interestingly, significant increases in the osteoid thickness, osteoid volume per bone volume, and the osteoid maturation time suggest that a mineralization defect occurs in mice unable to oxidize fatty acids obtained not only from BMAds, but also from circulation (Kim et al., 2017).

Given extensive literature describing interactions between BMAds and hematopoietic cells (Lee, Al-Sharea, Dragoljevic, & Murphy, 2018; Valet et al., 2020), we were surprised at the lack of substantial changes in hematopoietic progenitors, white blood cells, and red blood cells in BMAd-*Pnpla2*^-/-^ mice under basal and CR conditions. One possibility is that skeletal sites containing low BMAd numbers, including vertebrae, sternum, ribs and pelvis, may allow compensatory formation of circulating mature blood cells (Kricun, 1985). In addition, hematopoietic progenitors and mature blood cell populations were not substantially altered in bone marrow of long bones, with the exception of myeloid differentiation into neutrophils, suggesting that hematopoietic cellularity is generally maintained despite expansion of BMAT volume. Although hematopoietic progenitors and mature cells can use lipids for energy, glucose serve as the major source of energy for glycolysis and oxidative metabolism in these cell populations (Jeon, Hong, Kim, & Lee, 2020; Roy, Biswas, Verfaillie, & Khurana, 2018). It is perhaps unsurprising, given the importance of blood cell production in maintaining life, that there is a great deal of metabolic flexibility when it comes to fuel sources for hematopoiesis. While HSPC numbers are unaltered in our BMAd-*Pnpla2^-/-^* mice, myeloid lineage cell recovery is significantly blunted by CR and deficiency of BMAd-lipolysis following sublethal irradiation and in *in vitro* myeloid CFU assays. Myeloid progenitors from CR BMAd-*Pnpla2*^-/-^ mice had an impaired capacity to expand, suggesting that progenitors were reprogrammed in response to energy deficiency in the bone marrow niche. We did not profile the metabolic changes of HSPCs in CR BMAd-*Pnpla2*^-/-^ mice, but it has been reported that fatty acid oxidation is required for HSC asymmetric division to retain the stem cell properties (Ito et al., 2012).

In summary, we have developed a novel mouse model to specifically evaluate the importance of BMAds as a local energy source. We report that BMAds are a local energy source that support myeloid cell lineage regeneration following irradiation, and maintain progenitor differentiation capacity when systemic energy is limiting. In addition, we find that BMAT serves a highly specialized function to maintain bone mass and osteoblast function in times of elevated local energy needs such as with bone regeneration, increased whole body energy needs from cold exposure, or when dietary energy is limited due to CR.

### Limitations of Study

There are some limitations to the BMAd-Cre mice that should be noted. For instance, expression of Cre from the IRES within the 3’-UTR of *Adipoq* is relatively low, and thus rates of recombination are less frequent in young mice than optimal. Mice that are older than 16 weeks are suggested for future usage of this Cre mouse model. However, bone formation and turnover rates decrease with age both in mice and humans (Fatayerji & Eastell, 1999; Ferguson, Ayers, Bateman, & Simske, 2003), which may provide challenges for mechanistic studies. The IRES-Cre cassette also causes hypoadiponectinemia so experimental design must include appropriate controls with BMAd-Cre positive mice with or without floxed genes. Although a total absence of adiponectin reduces bone mass (Naot et al., 2016; Yang et al., 2019), mice with hypoadiponectinemia described here did not exhibit metabolic or bone phenotypes. In retrospect, a better approach might have been to accept loss of one *Adipoq* allele and insert the FAC cassette at the start site of the adiponectin coding region to promote high levels of Cre expression. This strategy might also have allowed inducible knockout of BMAd genes with tamoxifen through expression of CreERT2, which we found to be too inefficient to be functional when inserted into the 3’-UTR of *Adipoq*, with only 5 to 8% BMAds labeled in tamoxifen-treated adult mice. Of note, all mice used in these studies were on a mixed SJL and C57BL/6J background, and mouse strain influences bone mass, and responsiveness of bone to stressors.

## Materials and Methods

### KEY RESOURCES TABLE

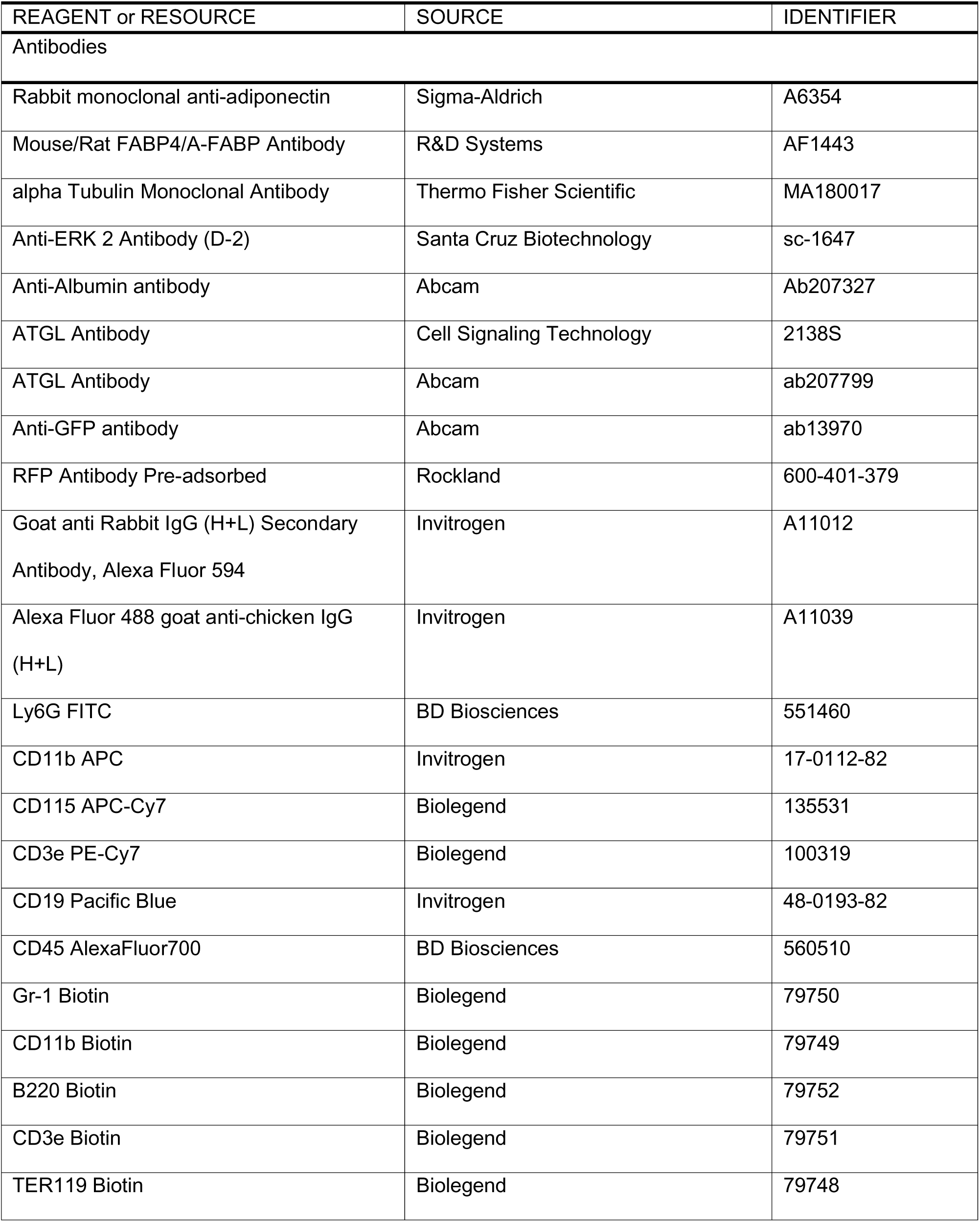
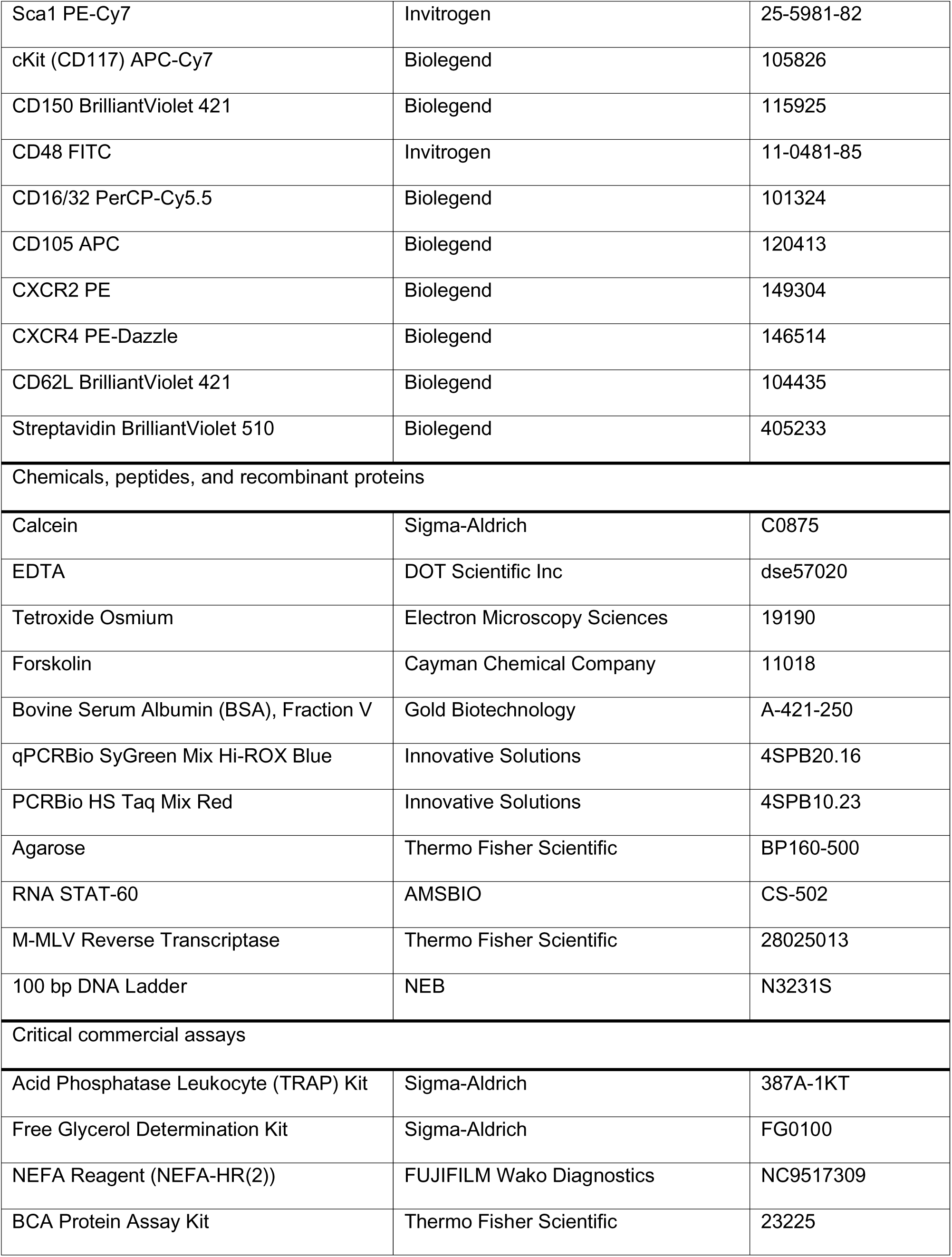
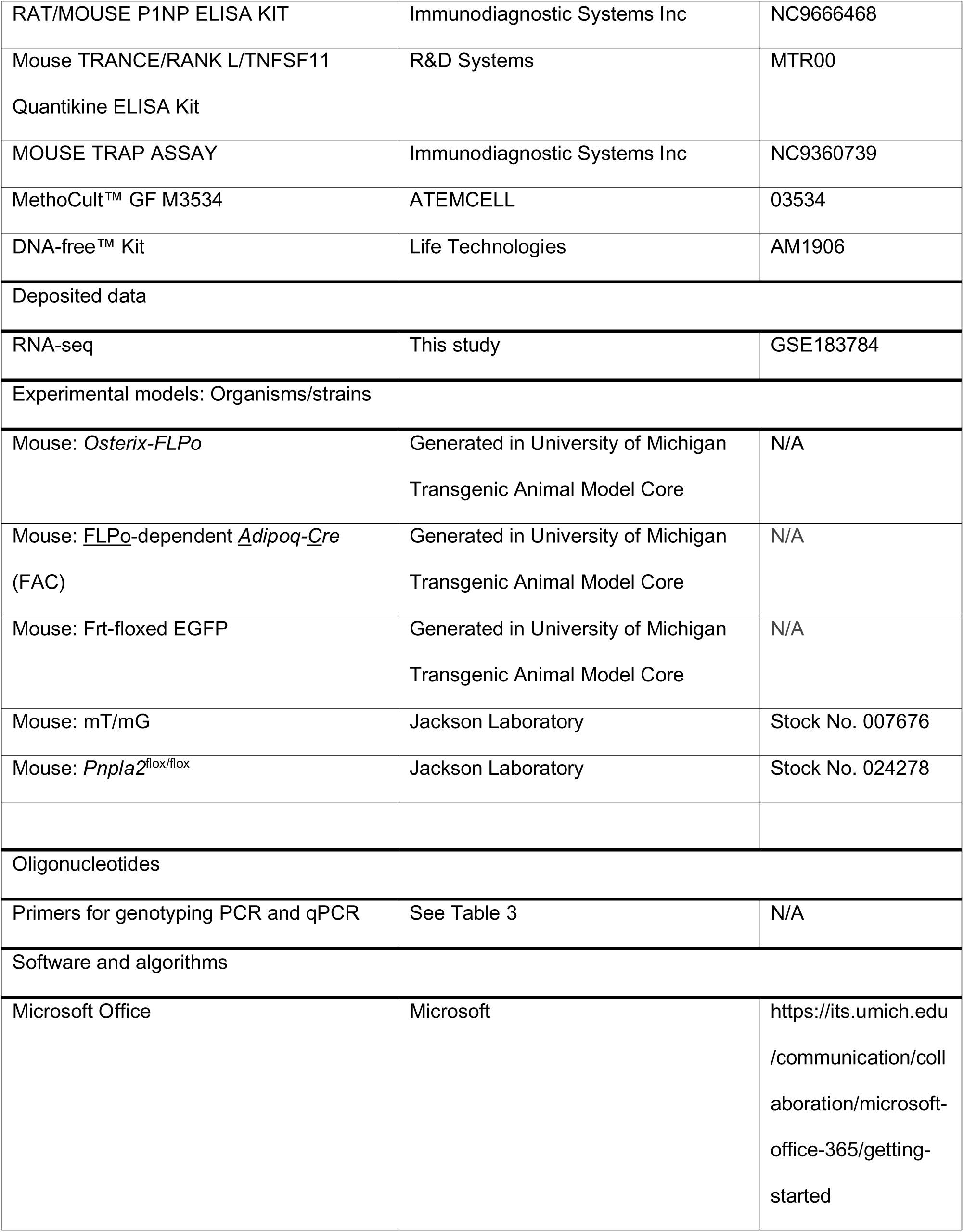
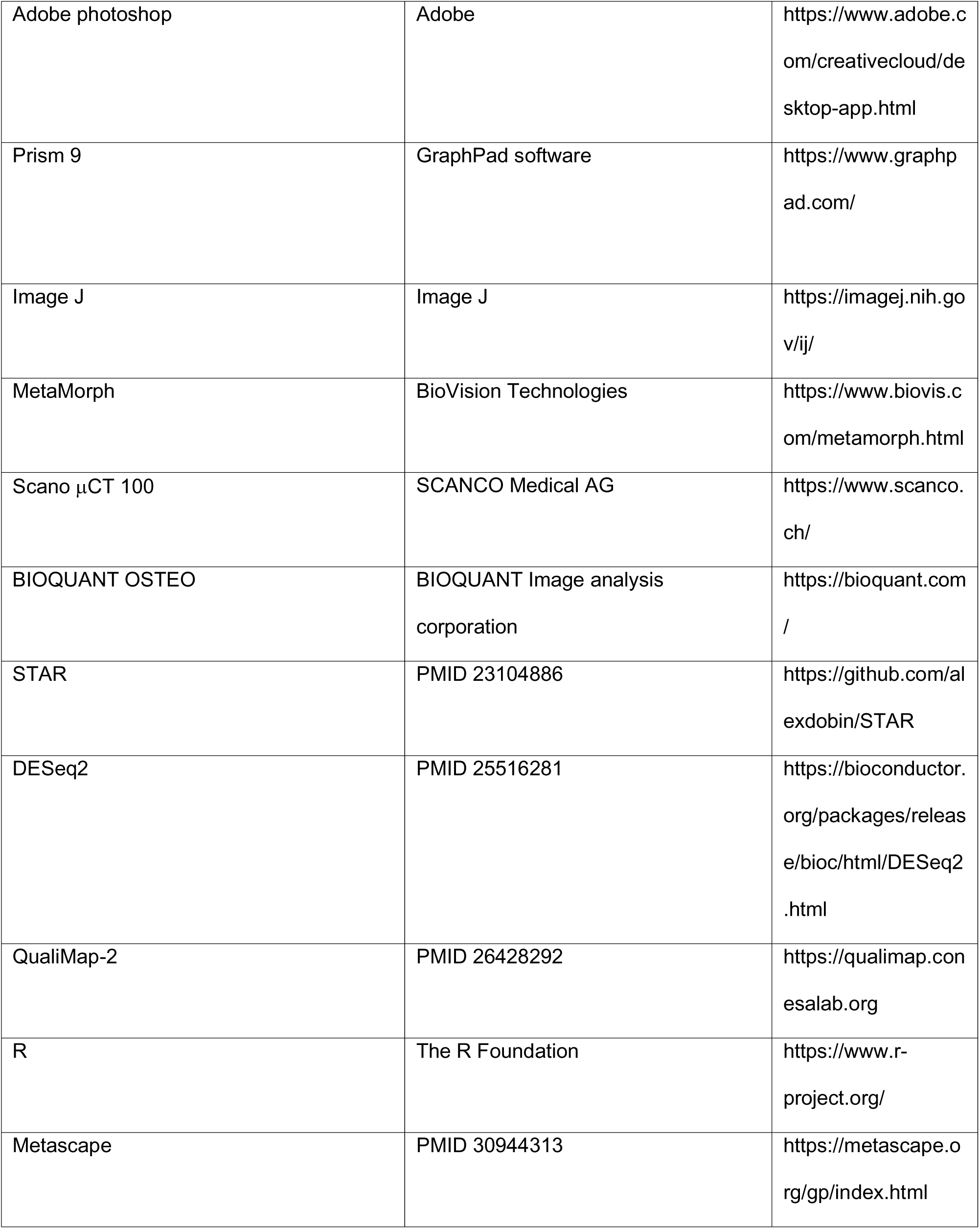
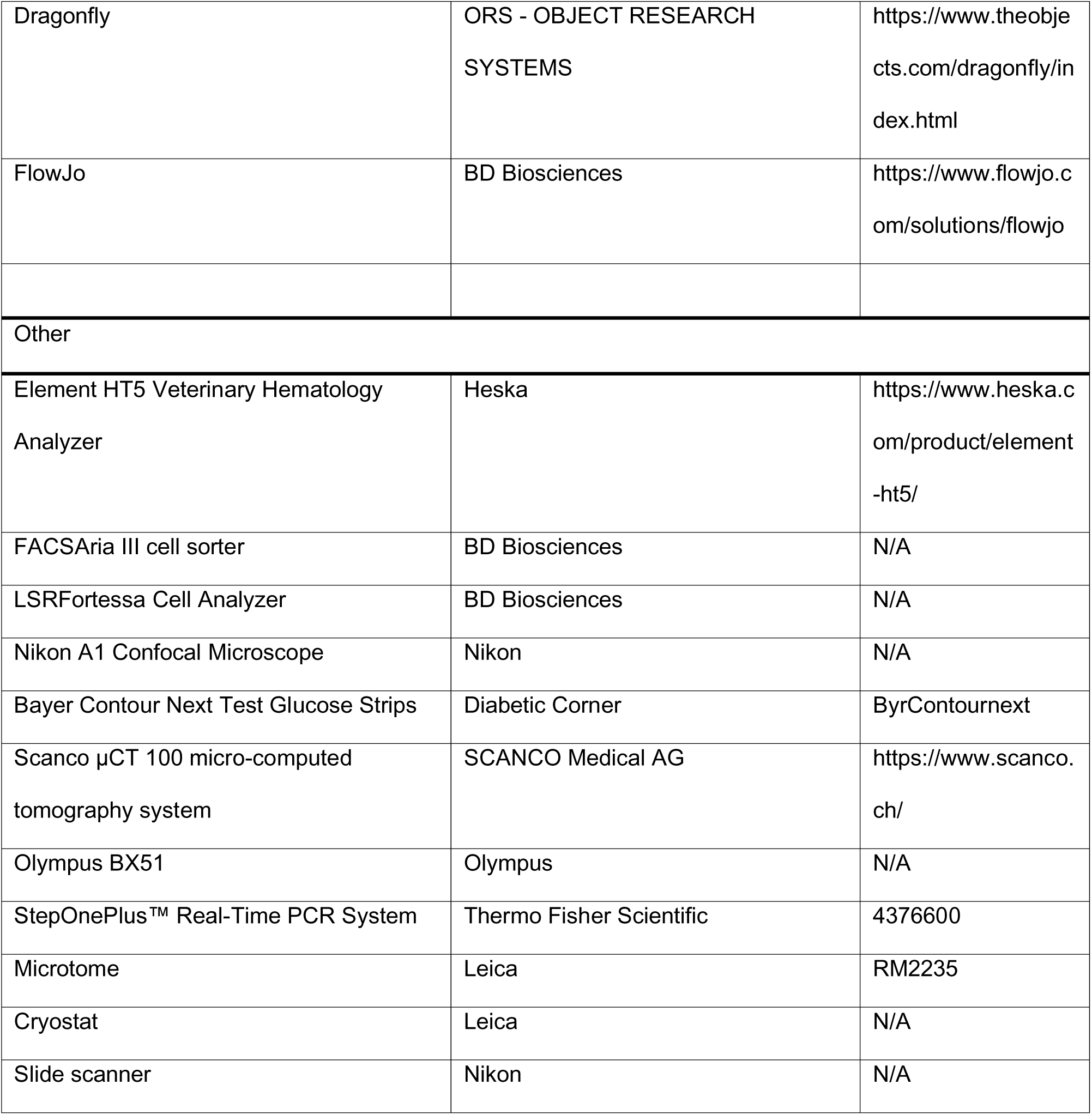

### Resource Availability

#### Lead contact

Further information and requests for resources and reagents should be directed to and will be fulfilled by the Lead Contact, Ormond A. MacDougald.

#### Materials availability

Our *Osterix-FLPo* and <u>FLPo</u>-dependent *<u>A</u>dipoq-<u>C</u>re* (FAC) mouse models will be available to investigators upon request. All the other data and materials that support the findings of this study are available within the article and supplemental information, or available from the authors upon request.

#### Data availability

The accession number for the BMAT bulk RNA seq data reported in this paper is GEO: GSE183784. This paper does not report original code.

### Animal

#### Generation of BMAd-specific Cre mouse model

To create a BMAd-specific Cre mouse model, we expressed mouse codon-optimized FLP (FLPo) (Raymond & Soriano, 2007) from the *Osterix* (*Sp7*) locus to recombine and activate a Cre expressed from the *Adipoq* locus. After designing sgRNA target sequences against 3’-UTR of endogenous *Osterix* (sgRNA: gatctgagctgggtagaggaagg) and *Adipoq* (sgRNA1: tgaacaagtgagtacacgtgtgg; sgRNA2: cagtgagcagaaaaatagcatgg) genes with the prediction algorithm available at http://crispor.tefor.net, we cloned sgRNA sequences into an expression plasmid bearing both sgRNA scaffold backbone (BB) and Cas9, pSpCas9(BB) (Ran et al., 2013), which is also known as pX330 (Addgene plasmid ID: 42230; Watertown, MA). Modified pX330 plasmids were injected into fertilized ova. After culture to the blastocyst stage, Cas9 activity was confirmed by sequencing the predicted cut site. We then inserted a DNA fragment containing an in-frame fusion protein between endogenous *Osterix* and FLPo, separated by coding sequence for P2A (porcine teschoviral-1) self-cleavage site to allow the full-length proteins to function independently. We also inserted into the 3’-UTR of *Adipoq* an <u>IRES-F3-Frt-reversed Cre-F3-Frt</u> cassette with ∼1kb of 5’ and 3’ flanking homology-arm sequence. The targeting vectors (Cas9 expression plasmid: 5 ng/ul; donor DNA: 10 ng/ul) were injected into fertilized mouse eggs, which were transferred into pseudopregnant recipients to obtain pups. Tail DNA was obtained for genomic PCR and Sanger sequencing to screen F0 founders for desired genetic modifications at the expected locations. These F0 generation chimeric mice were mated with normal mice to obtain mice that are derived exclusively from the modified fertilized eggs. Germline transmitted mice (F1 generation) carrying the designated genetic modifications were bred with reporter mice to validate the activity and efficiency of FLPo and Cre recombinase. These mice were generated by University of Michigan Transgenic Animal Model Core and MDRC Molecular Genetics Core.

#### Validation of Osterix-FLPo specificity and efficiency

*Osterix*-*FLPo* mice were bred with FLP-dependent EGFP reporter mice (derived from https://www.jax.org/strain/012429; Bar Harbor, ME), provided by Dr. David Olson from the University of Michigan MDRC Molecular Genetics Core. Fresh tissue confocal was performed to validate EGFP expression.

#### Validation of the specificity and efficiency of FLPo-activated Adipoq-Cre (FAC) mice

Among 80 F0 generation pups, 13 mice carried the FAC cassette. These founder mice were bred with *Osterix*-*FLPo* mice and mT/mG reporter mice (Stock No. 007676, Jackson Laboratory; Bar Harbor, ME) to further validate Cre recombinase activity and specificity via fresh tissue confocal and immunofluorescent histology. Every single mouse in F1 generation was confirmed and separated for future breeding. We finally selected F0-#693→ F1-#4376 for BMAd-specific knockout lines. Of note, *Osterix-FLPo* and *FAC* mice were generated on a mixed SJL and C57BL/6J background. *Pnpla2*^flox/flox^ mice purchased from Jackson laboratory (Stock No. 024278; Bar Harbor, ME) were on a C57BL/6J background.

#### Animal housing

Mice were housed in a 12 h light/dark cycle in the Unit for Laboratory Animal Medicine at the University of Michigan, with free access to water. Mice were fed *ad libitum* or underwent caloric restriction, as indicated. All procedures were approved by the University of Michigan Committee on the Use and Care of Animals with the protocol number as PRO00009687.

#### Animal Procedures

1. 30% CR. After acclimation to single-housing for 2 weeks, and control diet (D17110202; Research Diets; New Brunswick, NJ) for a week, daily food intake was measured for another week. Mice were then fed a 30% CR diet (D19051601; Research Diets; New Brunswick, NJ) daily at ∼6 pm, prior to onset of the dark cycle.
2. Ovariectomy. Female mice at 16 weeks of age underwent ovariectomy (OVX). After 2 weeks recovery, they were either fed an *ad libitum* or 30% CR diet for 12 weeks.
3. Proximal tibial defect. Surgeries were performed under isoflurane anesthesia, and subcutaneous 0.1 mg/kg buprenorphine was given in 12-h intervals for peri-/post-operative pain management. The proximal tibial defects were obtained by drilling a hole through anterior cortical and trabecular bone, 1 to 2 mm below the epiphyseal growth plate, with a 0.7 mm low-speed drill.
4. Cold exposure. Mice were single-housed without nesting materials in thermal chambers for 3 weeks at 5°C.
5. Glucose Tolerance Test (GTT). Mice were fasted overnight (16-18 hours). Body weight and fasting glucose levels were measured, followed by an intraperitoneal (i.p.) injection of glucose (1g glucose/kg body weight). Blood glucose was measured with Bayer Contour test strips at 15, 30, 60, 90 and 120 min time points by cutting the tip of tails.
6. Whole body irradiation. Irradiations were carried out using a Kimtron IC320 (Kimtron Medical; Oxford, CT) at a dose rate of ∼4 Gy/minute with total dosage of 6 Gy in the University of Michigan Rogel Cancer Center Experimental Irradiation Shared Resource. Dosimetry was carried out using an ionization chamber connected to an electrometer system that is directly traceable to a National Institute of Standards and Technology calibration.

##### Fresh tissue confocal microscopy

*Osterix*-driven EGFP expression and BMAd-Cre-driven mT/mG reporter mice were sacrificed. Fresh tissues were collected immediately and put in ice-cold PBS, which was protected from light. Soft tissues including adipose tissues and liver were cut into small pieces and placed in a chamber for confocal imaging (Nikon Ti-E Inverted Microscope; Minato City, Tokyo, Japan). Bones were bisected, and butterflied on a coverslip for imaging. Both white field and fluorescent images were taken under 200x magnification.

##### Histology and histomorphometry

Tissue histology was performed essentially as described previously (Z. Li et al., 2019). Briefly, soft tissues were fixed in formalin, and embedded in paraffin for sectioning. Tibiae were fixed in paraformaldehyde, decalcified in EDTA for at least 2 weeks, and followed by post-decalcification fixation with 4% paraformaldehyde. Bone tissues were then embedded in paraffin, sectioned. After staining with hematoxylin and eosin (H&E), soft tissues were imaged with an Olympus BX51 microscope. Bones were stained with H&E or Tartrate-Resistant Acid Phosphatase (TRAP; Sigma-Aldrich, MO) as indicated, and slides were scanned at 200X magnification. Static measurements include bone volume fraction, trabecular bone microstructure parameters, osteoblast number and surface, and osteoclast number and eroded surface. Undecalcified tibia was used for plastic sectioning. Mineralized trabecular bone and osteoid were evaluated with Goldner’s Trichrome Staining. For dynamic studies, calcein (C0857; Sigma-Aldrich, MO) dissolved in 0.02 g/ml sodium bicarbonate with 0.9% saline at 20 mg/kg was injected intraperitoneally nine- and two-days before sacrifice for quantification of mineral surface (MS), inter-label width (Ir. L. Wi), and mineral apposition rate (MAR) in tibia. Calculations were made with Bioquant Osteo 2014 (Nashville, TN) software in a blinded manner (Merceron et al., 2014; Morse et al., 2014).

##### μCT analysis

Tibiae were placed in a 19 mm diameter specimen holder and scanned over the entire length of the tibiae using a μCT system (μCT100 Scanco Medical, Bassersdorf, Switzerland). Scan settings were: voxel size 12 µm, 70 kVp, 114 µA, 0.5 mm AL filter, and integration time 500 ms. Density measurements were calibrated to the manufacturer’s hydroxyapatite phantom. Analysis was performed using the manufacturer’s evaluation software with a threshold of 180 for trabecular bone and 280 for cortical bone.

##### Marrow fat quantification by osmium tetroxide staining and μCT

After analyses of bone variables, mouse tibiae were decalcified for osmium tetroxide staining, using our previously published method (Scheller et al., 2015). In addition, a lower threshold (300 grey-scale units) was used for proximal tibial rBMAT quantification because density of osmium staining is low due to smaller adipocyte size, and with threshold as 400 grey-scale units for cBMAT in distal tibia.

##### Immunofluorescent staining

Decalcified tibiae were embedded in OCT compound and used for frozen sectioning at 15 μm. Excess OCT was removed, and tibial tissues were blocked with 10% goat serum for one hour at room temperature. Primary antibodies for GFP (1:500) and RFP (1:200; Figure 1F) or ATGL (1:100; Figure 2B) were then added to slides and incubated with bone tissues overnight at 4°C. Secondary antibodies (goat anti Rabbit IgG, Alexa Fluor 594 & goat anti-chicken IgG Alexa Fluor 488 or donkey anti rabbit IgG (H+L), Alexa Fluor 488) were added to slides following three washes. Two hours later, DAPI staining was performed. Slides were mounted with prolong gold antifade reagent and imaged.

##### RNA extraction and quantitative real-time PCR (qPCR)

RNA was extracted from tissues after powdering in liquid nitrogen and lysis in RNAStat60 reagent in a pre-cooled dounce homogenizer. Quantitative PCR was performed using an Applied Biosystems QuantStudio 3 qPCR machine (Waltham, MA). Gene expression was calculated based on a cDNA standard curve within each plate and normalized to expression of the geometric mean of housekeeping genes *Hprt, Rpl32A* and *Tbp*.

##### Immunoblot

Detection of proteins by immunoblot was as described previously (Mori et al., 2021).

##### *Ex vivo* lipolysis

Distal tibial BMAT plugs were flushed out from the bone, and four plugs were pooled into one well of a 96-well plate for each n. Pre-warmed 2% BSA HBSS was added to BMAT explants with vehicle (DMSO) or forskolin (FSK, 5 μM), to activate lipolysis (Litosch, Hudson, Mills, Li, & Fain, 1982). Cultured media was collected hourly for 4 hours, and glycerol and NEFA concentrations were measured with commercially available kits as listed above in reagents.

##### CFU assay

Bone marrow cells of BMAd-*Pnpla2*^+/+^ and BMAd-*Pnpla2*^-/^ mice were obtained from femur and tibia. To isolate bone marrow, the bones were flushed with IMDM (Gibco 12440; Waltham, MA) containing penicillin-streptomycin antibiotic (Gibco 15270-063; Waltham, MA). Pelleted cells were counted with a hemocytometer, 1X10^4^ cells were plated in Methocult medium (Stem cell M3534; Vancouver, Canada) in 35 mm culture dishes, and cells were then incubated at 37°C in 5% CO_2_ with ≥ 95% humidity for seven days. CFU-G, CFU-M and CFU-GM colonies on each plate were counted using a microscope with a 4X objective lense.

##### Bulk RNA sequencing

Distal tibial plugs (cBMAT) from two animals were pooled together for each bulk RNAseq sample. Total RNA was isolated from BMAT for strand specific mRNA sequencing (Beijing Genomics Institute, China). Over twenty million reads were obtained using a paired-end 100 bp module on DNBSEQ platform. The quality of the raw reads data was checked using FastQC (v.0.11.9) and the filtered reads were aligned to reference genome (UCSC mm10) using STAR with default parameters. All samples passed the post-alignment quality check (QualiMap (v.2.2.1). The DEseq2 method was used for differential expression analysis with genotype (*Pnpla2*^+/+^ vs. *Pnpla2*^-/-^) and treatment (CR vs. *ad libitum*) as the main effects. Gene ontology analysis was done using MetaScape. The Principal Component Analysis (PCA) plot was generated using the PlotPCA function building in DESeq2 package. To compare the number of differentially expressed genes between genotype/treatment groups, volcano plots were constructed using the Enhanced Volcano package under R environment. Heatmap plots were generated using pheatmap package under R environment and the complete-linkage clustering method was used for the hierarchical cluster of genes. These data are available through NCBI GEO with the following accession number, GSE183784.

##### Flow cytometry

Femurs were isolated from mice. Bone marrow was harvested by flushing the femurs with 1 mL of ice-cold PEB (1X PBS with 2 mM EDTA and 0.5% bovine serum albumin). Red blood cells were lysed once by adding 1 mL of RBC Lysis Buffer (155 mM NH_4_Cl, 10 mM KHCO_3_, 0.1 mM EDTA) and gently pipetting to mix. Cells were immediately pelleted by centrifugation and resuspended in 1 mL of ice-cold PEB. Cells were stained for 30 minutes in PEB buffer with the indicated antibodies below and analyzed on the BD LSRFortessa or BD FACSAria III. Data was analyzed using FlowJo software (BD Biosciences, version 10.8). Dead cells and doublets were excluded based on FSC and SSC distribution. To stain for mature leukocytes antibodies used were against CD45, Ly6G, CD11b, CD115, CD19, and CD3e. All CD45^+^ cells were gated first for further identification. Neutrophils were defined as Ly6G^+^CD11b^+^, monocytes were defined as Ly6G^-^ CD11b^+^CD115^+^, B cells were defined as Ly6G^-^CD11b^-^CD19^+^, and T cells were defined as Ly6G^-^CD11b^-^CD3e^+^. To stain for bone marrow neutrophil populations, antibodies used were against Ly6G, CD11b, cKit, CXCR2, CXCR4, and CD62L. Pre-neutrophils were defined as Ly6G^+^CD11b^+^cKit^+^, immature neutrophils were defined as Ly6G^+^CD11b^+^cKit^-^CXCR2^lo^, and mature neutrophils were defined as Ly6G^+^CD11b^+^cKit^-^CXCR2^hi^. To stain for hematopoietic stem and progenitor cells (HSPCs) antibodies used were against a lineage panel (Gr-1, CD11b, B220, CD3e, TER119), cKit, Sca-1, CD150, CD48, CD105, and CD16/32. After gating on the lineage^-^ population, HSPCs were defined as follows; HSCs as LSKCD150^+^CD48^-^, MPPs as LSKCD150^-^CD48^-^, HPC1 as LSKCD150^-^CD48^+^, HPC2 as LSKCD150^+^CD48^+^, GMPs as LKCD150^-^CD16/32^+^, PreGMs as LKCD150^-^CD105^-^, PreMegEs as LKCD150^+^CD105^-^, and PreCFUe as LKCD150^+^CD105^+^. LSK = lineage^-^Sca1^+^cKit^+^, LK = lineage^-^cKit^+^.

##### Bone regeneration analysis

Nine days after the proximal tibial defect surgery, proximal tibiae were collected for μCT scanning. After construction, new generated trabecular and cortical bone were quantified by Dragonfly software (Montréal, Canada). Under the full view of 3D and 2D images, a cylinder-shaped region of interest (ROI) was defined in trabecular or cortical bone defect area (as shown in Figure 5A) with 3D dimension as 0.3 mm (diameter) x 0.7 mm (height) for trabecular bone and 0.3 mm (diameter) x 0.2 mm (height) for cortical bone. An automatic split at otsu threshold for each bone was collect first, and then mean threshold for a whole cohort was calculated. This average threshold was applied to each bone to normalize bone volume fraction and mineral content.

##### Statistics

We calculated the minimal animal number required for studies based on the mean and SD values to make sure we had adequate animals per group to address our hypothesis. All the mice were randomly assigned to the indicated groups. Although the investigators responsible for group allocation were not blinded to the allocation scheme, they were blinded to group allocation during data collection, and the investigators responsible for analyses were blinded to the allocation scheme.

Significant differences between groups were assessed using a two-sample *t*-test or ANOVA with post-tests as appropriate: one-way ANOVA with Tukey’s multiple comparisons test, two-way ANOVA with Sidak’s multiple comparisons test and three-way ANOVA analysis as appropriate. All analyses were conducted using the GraphPad Prism version 9. All graphical presentations are mean +/- SD. For statistical comparisons, a *P*-value of < 0.05 was considered significant. All experiments were repeated at least twice.

## Acknowledgements

This work was supported by grants or fellowships from the NIH to OAM (R01 DK62876; R24 DK092759; R01 DK126230; R01 AG069795), SMR (T32 GM835326; F31 DK12272301), DPB (T32 HD007505; T32 GM007863), KS (R01DK115583), KTL (T32 DK071212; F32 DK122654), RLS (T32 DK101357; F32 DK123887), KDH (R01AR066028), and CJR (R24DK092759). ZL was supported by a fellowship from the American Diabetes Association (1-18-PDF-087) and EB from the American Heart Association (20-PAF00361). This research was also supported by a Pilot & Feasibility grant and core facilities of the Michigan Integrative Musculoskeletal Health Core Center (P30 AR069620), Michigan Diabetes Research Center (P30 DK020572), Michigan Nutrition and Obesity Center (P30 DK089503), and University of Michigan Comprehensive Cancer Center (P30 CA046592).

## Author contributions

ZL, KS, KDH, CJR, and OAM conceived the studies and planned the experimental design. ZL, JH, JZ, EB, DPB, HM, KG, KTL, RLS, SMR, and SA performed the experiments. ZL, HY, and OAM analyzed the data. ZL and OAM wrote the manuscript, while all other authors edited and approved the final manuscript.

## Figures and figure supplement list

**Figure 1 - figure supplement 1.**
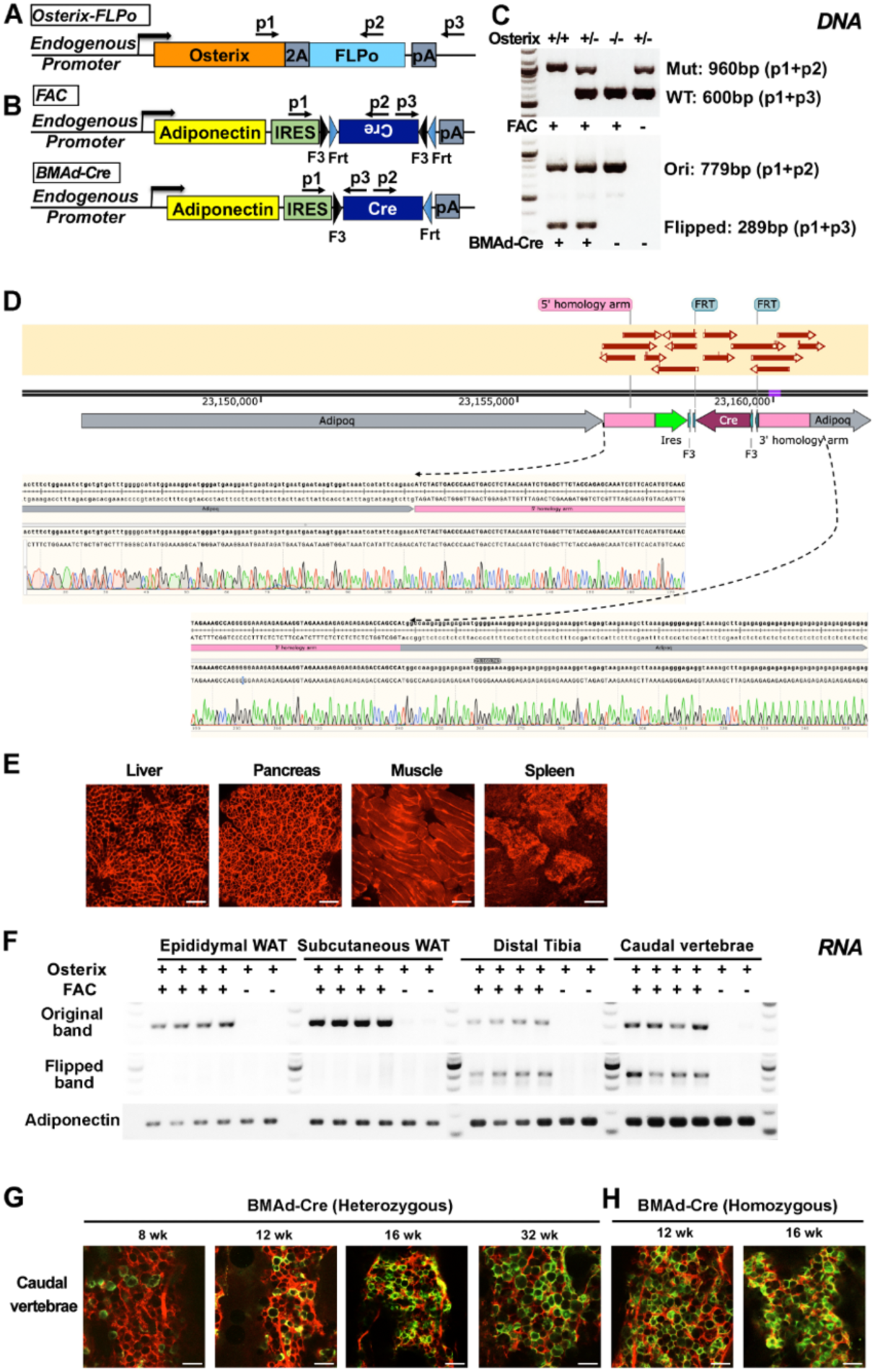
Genotyping strategy and validation of BMAd-Cre. A-C. Schematic of genotyping primer designing for *Osterix*-*FLPo* (A) and FLPo activated *Adipoq*-*Cre* (FAC; B), and representative genotyping results (C). Mut= Mutation; WT=wildtype; Ori=original band without conversion of reversed Cre cassette; Flipped=FLPo activated Cre (BMAd-Cre). D. Representative image of genomic PCR sequencing aligned with the endogenous allele. E. BMAd-Cre mice were bred with mT/mG reporter mice and the resulting BMAd-mT/mG mice were sacrificed at 16 weeks of age. Cellular fluorescence was evaluated by fresh tissue confocal microscopy. Scale bar; 100 μm. F. The flipped recombination band is found in mRNA of bone, but not WAT depots. Mice expressing *Osterix*-*FLPo* with or without FAC were sacrificed and mRNA was isolated. cDNAs from distal tibia, caudal vertebrae, epididymal and subcutaneous WATs were used for PCR and agarose gel electrophoresis. G, H. Efficiency of BMAd-Cre recombinase is age- and allele-dependent. Fresh caudal vertebrae collected from BMAd-mT/mG mice at indicated ages were bisected and used for confocal imaging. Scale bar; 100 μm.

**Figure 1 - figure supplement 2.**
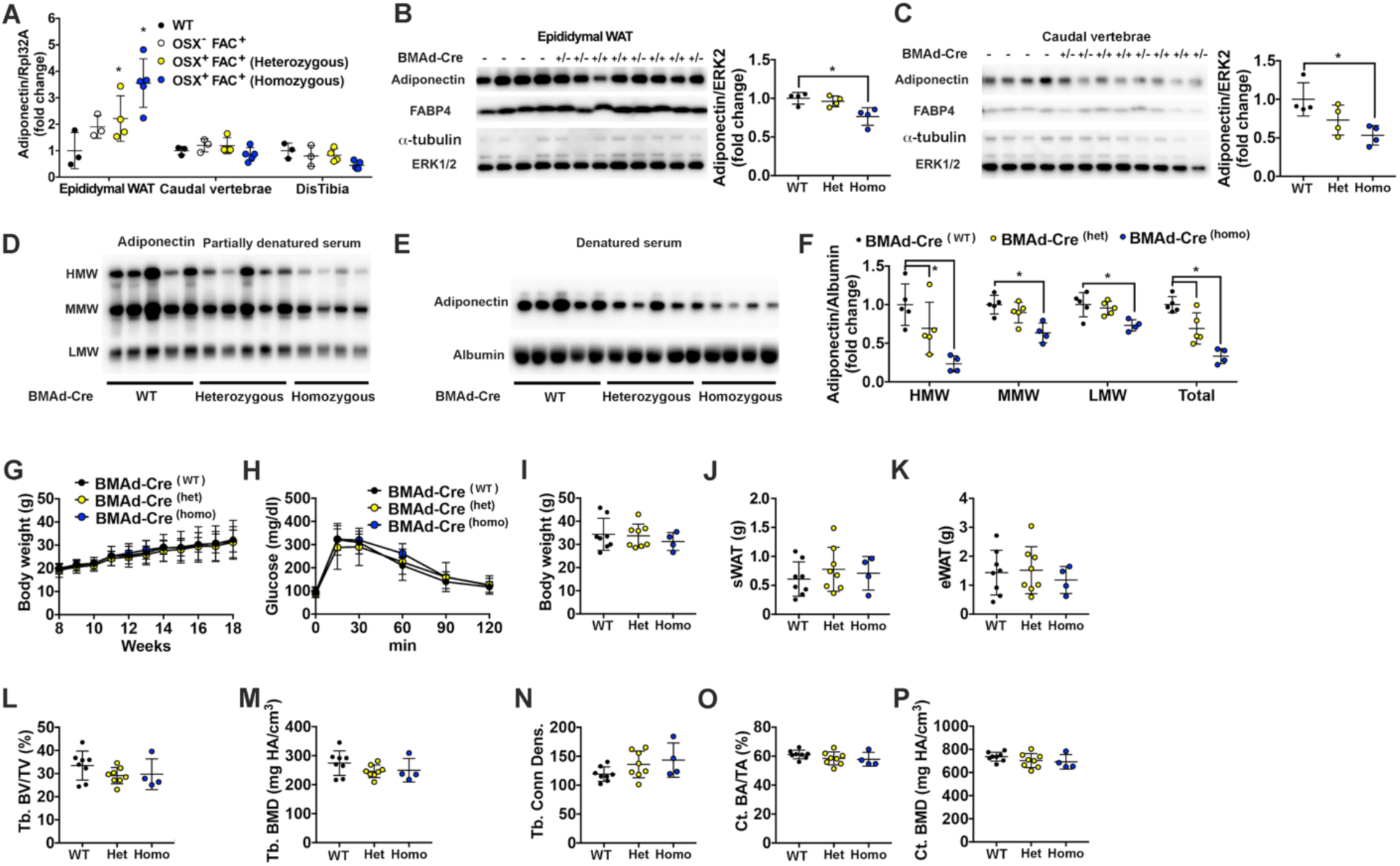
Insertion of IRES-Cre cassette in the 3’UTR of endogenous *Adipoq* decreases expression of adiponectin but does not cause a metabolic or bone phenotype. A-F. Endogenous adiponectin expression is decreased by IRES-Cre insertion. Mice expressing *Osterix*- *FLPo* with or without FAC were sacrificed. Epididymal WAT (eWAT), caudal vertebrae, distal tibiae and serum were collected. A. mRNA expression of adiponectin in eWAT, caudal vertebrae and distal tibiae from BMAd-Cre (OSX^+^FAC^+^) WT, Heterozygous, and Homozygous mice. B-C. Lysates from eWAT (B) and caudal vertebrae (C) were used for immunoblot analyses of adiponectin and FABP4, with *α*-tubulin and ERK1/2 as loading controls. Quantification was performed using Image J. D-F. Non-denatured (D) and denatured (E) serum from BMAd-Cre WT, heterozygous, or homozygous mice was used for immunoblot analyses of adiponectin, with albumin as a reference protein. High, medium and low molecular weight forms of adiponectin were quantified by Image J. G-P. Hypoadiponectinemia does not cause detectable phenotypes. Male BMAd-Cre WT, heterozygote, or homozygote mice were sacrificed at 18 weeks of age. Soft tissues and bones were collected. G-K. Body weight, glucose tolerance, and WAT depot weights are not changed in BMAd-Cre mice. L-P. Trabecular and cortical bone parameters were determined by μCT. Data are expressed as mean ± SD. * indicates *P* < 0.05 with one-way ANOVA with Tukey’s multiple comparisons test.

**Figure 3 - figure supplement 1.**
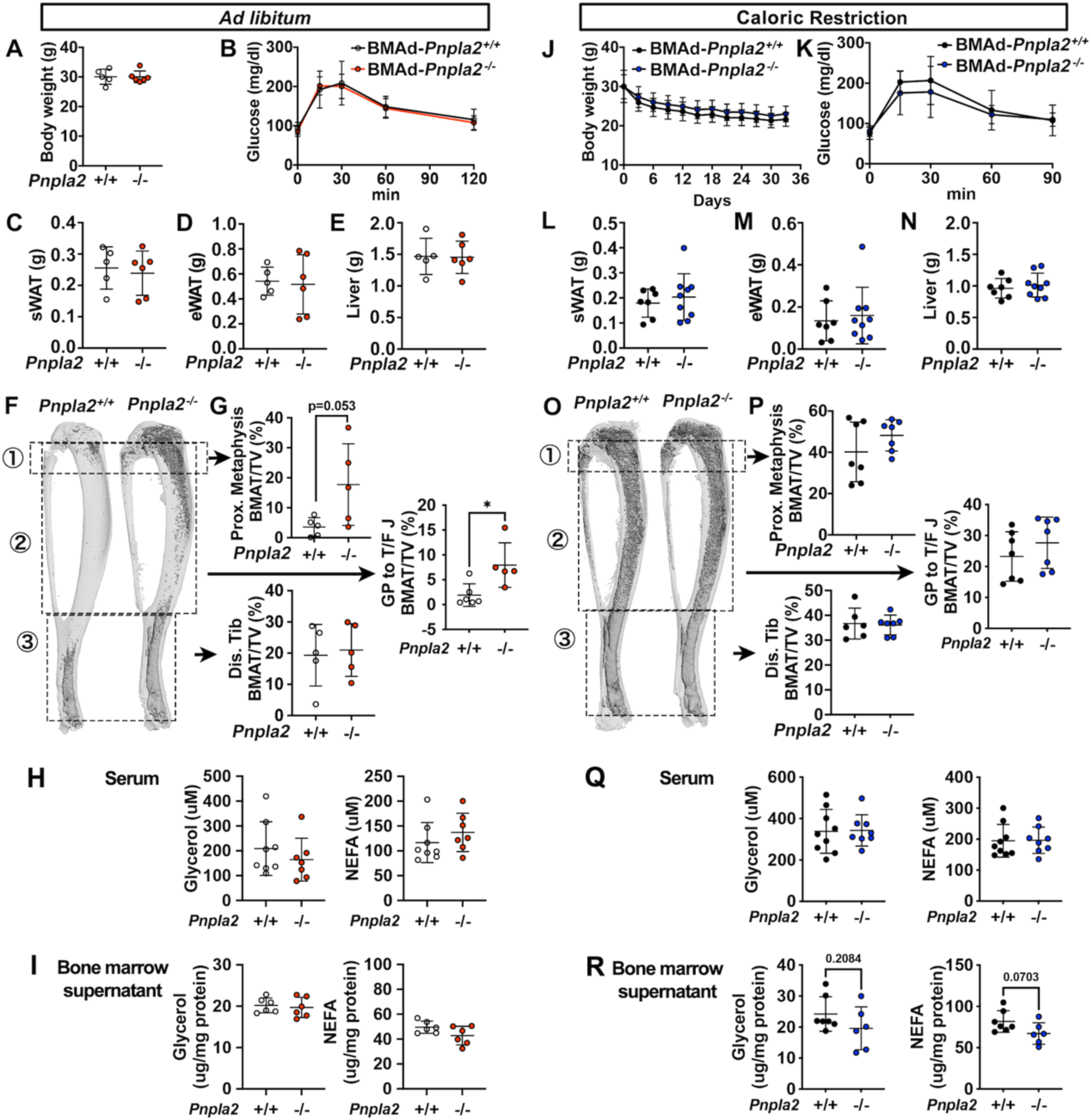
Blocking BMAd-lipolysis does not influence global metabolism when mice are fed *ad libitum* or calorically restricted. A-I. BMAd-*Pnpla2*^-/-^ male mice and littermate controls (*Pnpla2*^+/+^) were fed *ad libitum* until 24 weeks of age. Body weight (A), glucose tolerance test (B), and weights of subcutaneous WAT (sWAT), epididymal WAT (eWAT) and liver (C-E) were recorded. Decalcified tibiae were used for osmium tetroxide-staining and quantified by μCT analyses to measure the BMAT volume at proximal and distal ends, as indicated by boxed regions (F-G). Concentrations of glycerol and NEFA in serum (H) and bone marrow supernatant (I) were measured with colorimetric assay kits. Glycerol and NEFA contents in bone marrow supernatant were normalized to protein concentrations. J-R. BMAd-*Pnpla2*^-/-^ male mice and littermate controls (*Pnpla2*^+/+^) at 18 weeks of age and underwent a 30% CR for 6 weeks. Body weight changes (J) and glucose tolerance (K) were recorded. sWAT (L), eWAT (M), and liver (N) weights were measured during dissection. BMAT volume at proximal and distal tibia were quantified in osmium tetroxide-stained bones following μCT scanning (O-P). Concentrations of glycerol and NEFA in serum (Q) and bone marrow supernatant (R) were measured using colorimetric assays. Glycerol and NEFA contents in bone marrow supernatant were normalized to protein concentrations. Data are expressed as mean ± SD. * indicates *P* < 0.05 with a two-sample *t*-test.

**Figure 3 - figure supplement 2.**
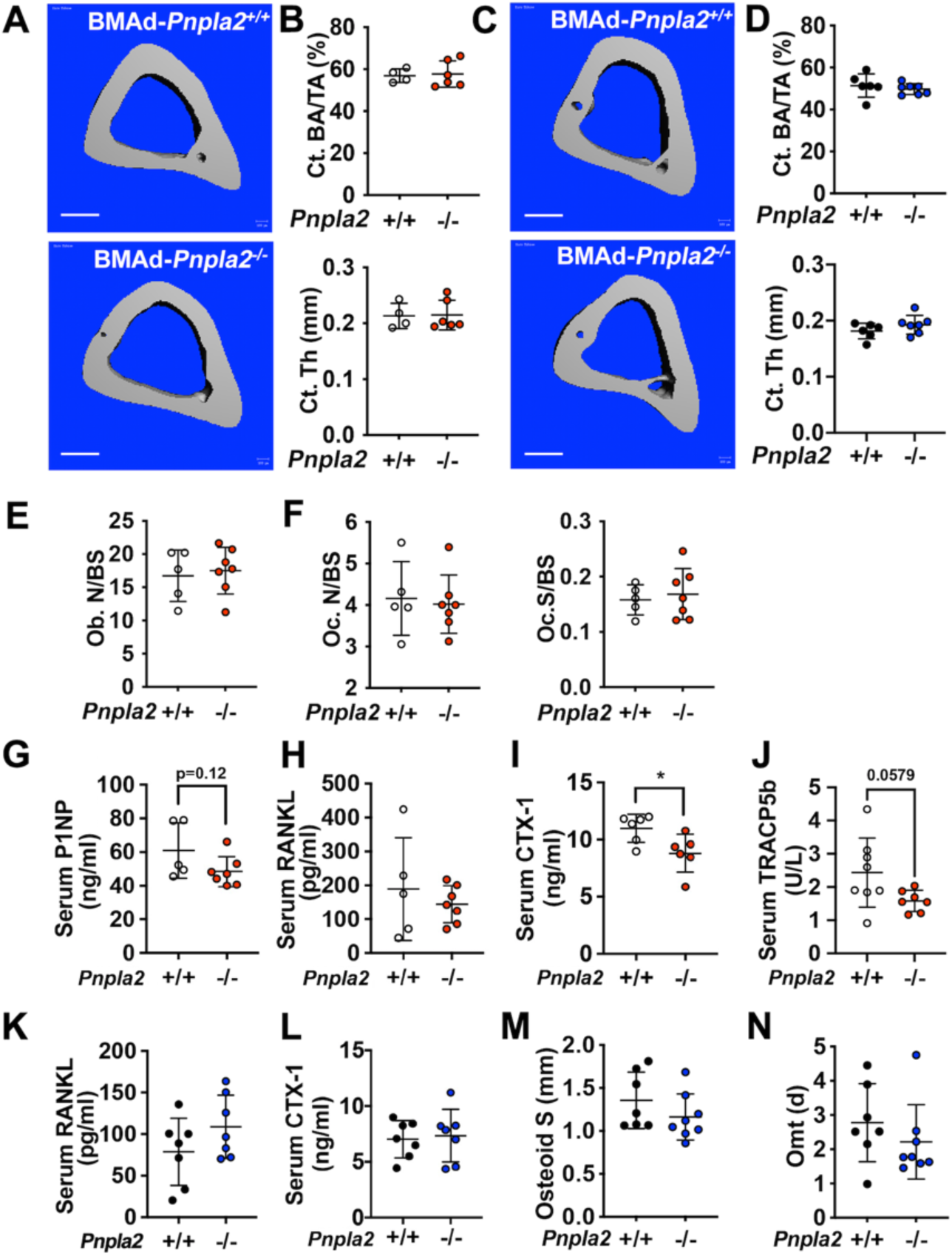
Cortical bone variables in BMAd-*Pnpla2*^-/-^ mice and other possible mechanisms for bone loss in BMAd-*Pnpla2*^-/-^ CR mice. A-D. Mouse tibiae from 24 weeks old *ad libitum* (A-B) or CR (C-D) mice were collected. Cortical bone area (CT. BA/TA) and thickness (Ct. Th) were measured by μCT. Scale bar; 500 μm. E-J. BMAd-*Pnpla2*^-/-^ male mice and littermate controls (*Pnpla2*^+/+^) were fed *ad libitum* until 24 weeks of age. E-F. Proximal tibial static histomorphometry was performed to calculate osteoblast number (Ob. N), osteoclast number (OC. N) and osteoclast surface (Oc. S) per bone surface (BS). G-J. Circulating P1NP, RANKL, CTX-1 and TRACP5b were measured with commercially available ELISA kits. K-N. BMAd-*Pnpla2*^-/-^ male mice and littermate controls (*Pnpla2*^+/+^) at 18 weeks of age underwent a 30% CR for 6 weeks. K-L. Circulating RANKL and CTX-1 were measured with commercially available ELISA kits. M-N. Histomorphometry analysis for osteoid surface and osteoid maturation time (Omt). Data are expressed as mean ± SD. * indicates *P* < 0.05 with a two-sample *t*-test.

**Figure 4 - figure supplement 1.**
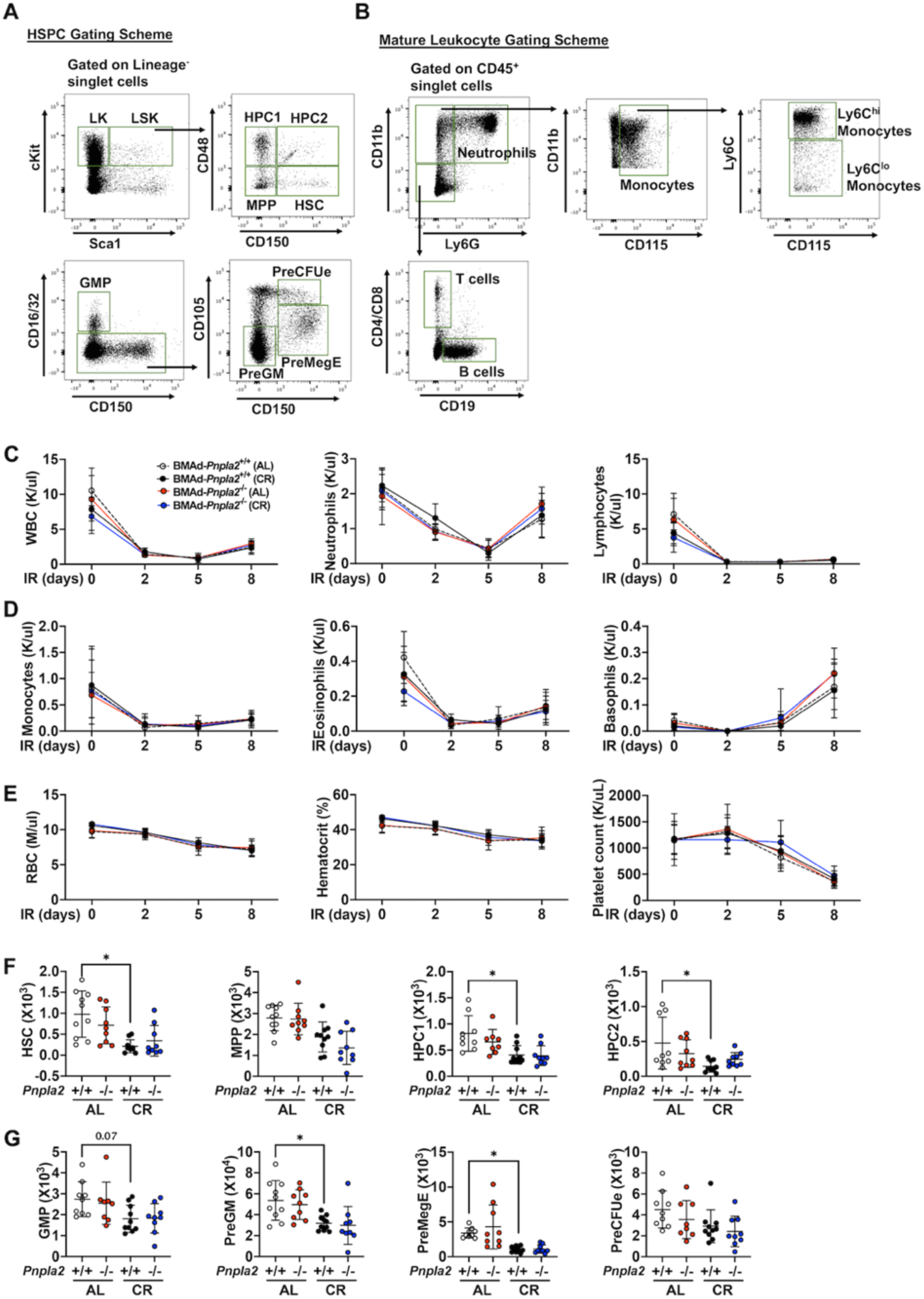
Flow cytometry strategies for hematopoietic cells and sublethal irradiation-induced hematopoietic regeneration in BMAd-*Pnpla2*^-/-^ mice. A-B. Hematopoietic cell gating strategies in hematopoietic stem/progenitor cells (HSPC, A) and mature leukocytes (B). C-G. BMAd-*Pnpla2*^-/-^ mice and littermate controls (*Pnpla2*^+/+^) underwent 30% caloric restriction (CR) for 20 weeks or remained on *ad libitum* (AL) diet, and then received a whole-body irradiation (6 Gy). Tail vein blood (∼50 ul) was collected every 2-3 days to monitor hematopoietic cell recovery. Mice were euthanized at day 9 post irradiation. C-E. Data from complete blood cell counts shows dynamic changes of white- and red blood cells before and after irradiation. F-G. Hematopoietic cells from two femurs were stained with cell markers to identify hematopoietic stem and progenitor cells. Data are expressed as mean ± SD. * indicates *P* < 0.05 with one-way ANOVA with Tukey’s multiple comparisons test.

**Figure 3 - figure supplement 3.**
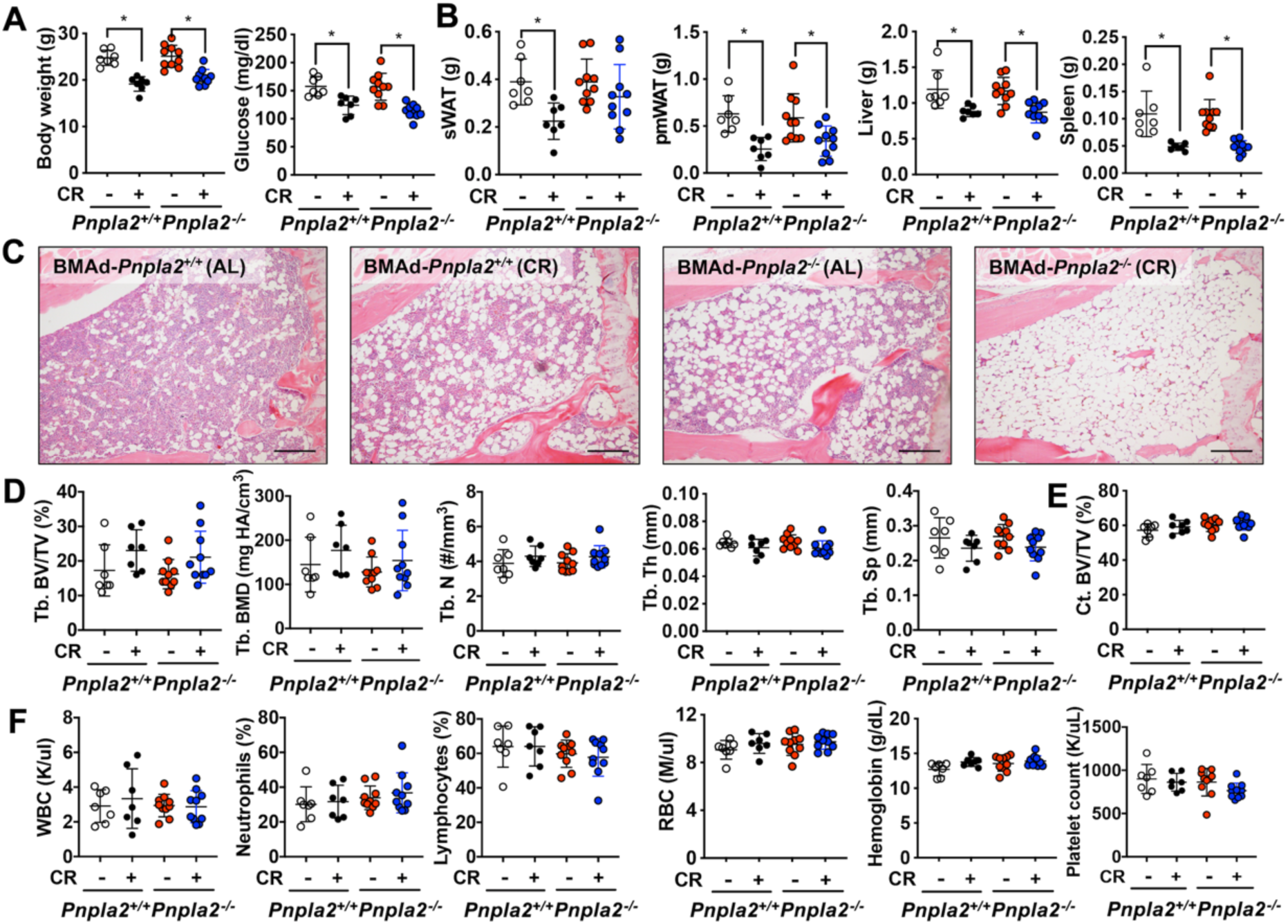
BMAd lipolysis is not required in female mice to maintain bone homeostasis under CR conditions. BMAd-*Pnpla2*^-/-^ female mice and their wildtype controls (*Pnpla2*^+/+^) at 18 weeks of age were fed a*d libitum* (AL) or underwent a 30% CR for another 6 weeks. A-B. Final body weight and random glucose levels were measured (A). sWAT, parametrial WAT (pmWAT), liver and spleen weights were recorded during dissection (B). C. Representative images from proximal tibiae were collected from decalcified and paraffin-sectioned bones. Scale bar; 200 μm. D-E. Trabecular and cortical bone variables were determined by μCT analysis. F. Complete blood counts (CBC) were performed to measure white- and red- blood cells in circulation. Data are expressed as mean ± SD. * indicates *P* < 0.05 with one-way ANOVA with Tukey’s multiple comparisons test.

**Figure 3 - figure supplement 4.**
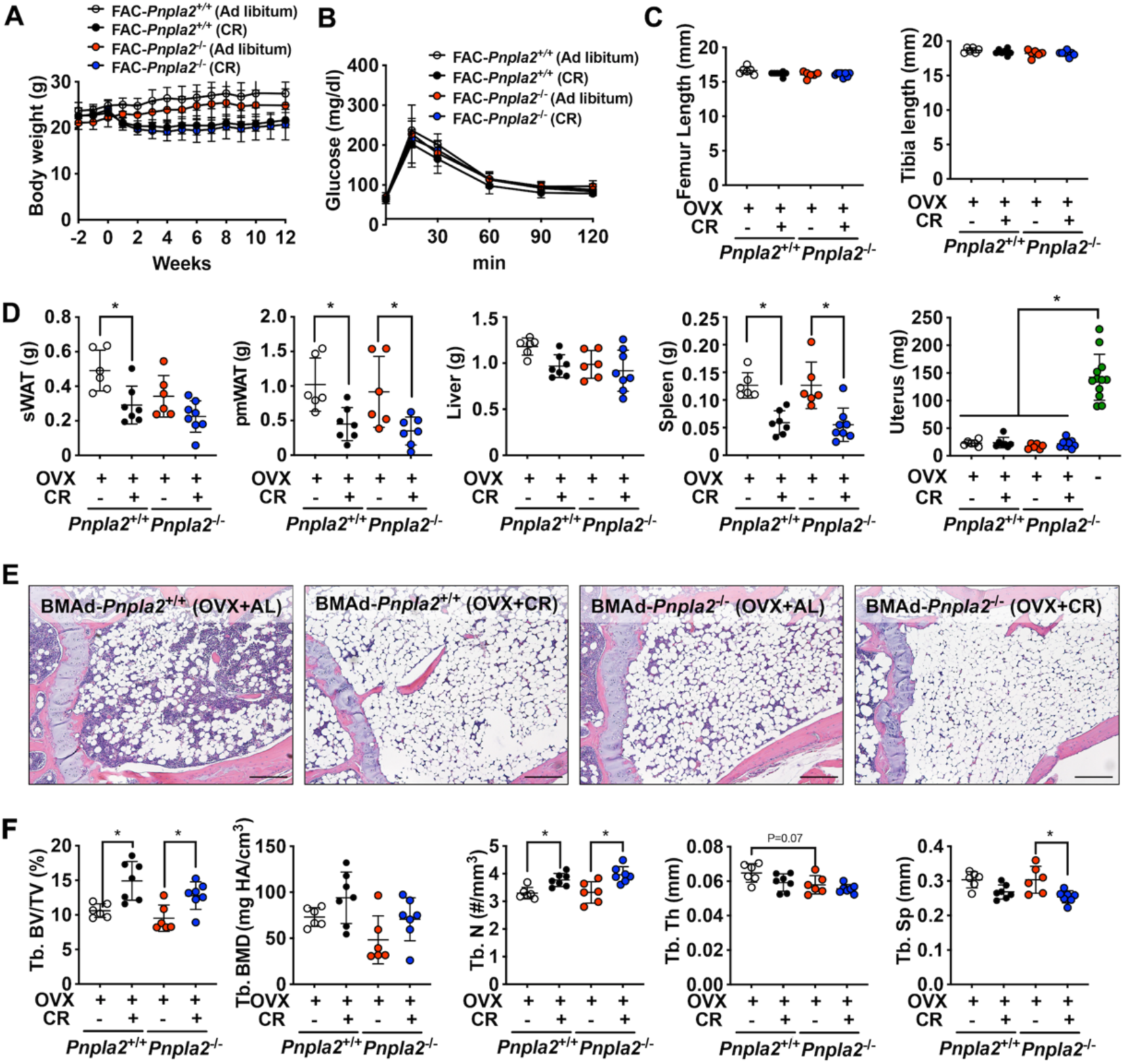
BMAd-lipolysis impairment in estrogen-deficient female mice does not affect CR-induced bone changes. 16 weeks old female mice underwent ovariectomy and recovered for 2 weeks, which were followed by 30% CR for 12 weeks. Changes in body weight (A) and glucose tolerance test (B) were recorded. Femoral and tibial lengths were measured (C). Weights of sWAT, pmWAT, liver, spleen, and uterus (green dots indicate sham mice) were measured during dissection (D). Trabecular bone parameters were determined by μCT (E). Tb.: trabecular bone; BV/TV: bone volume fraction; BMD: bone mineral density; N: number; Th: thickness; Sp: separation. Data are expressed as mean ± SD. * indicates *P* < 0.05 with one-way ANOVA with Tukey’s multiple comparisons test.

**Figure 5 - figure supplement 1.**
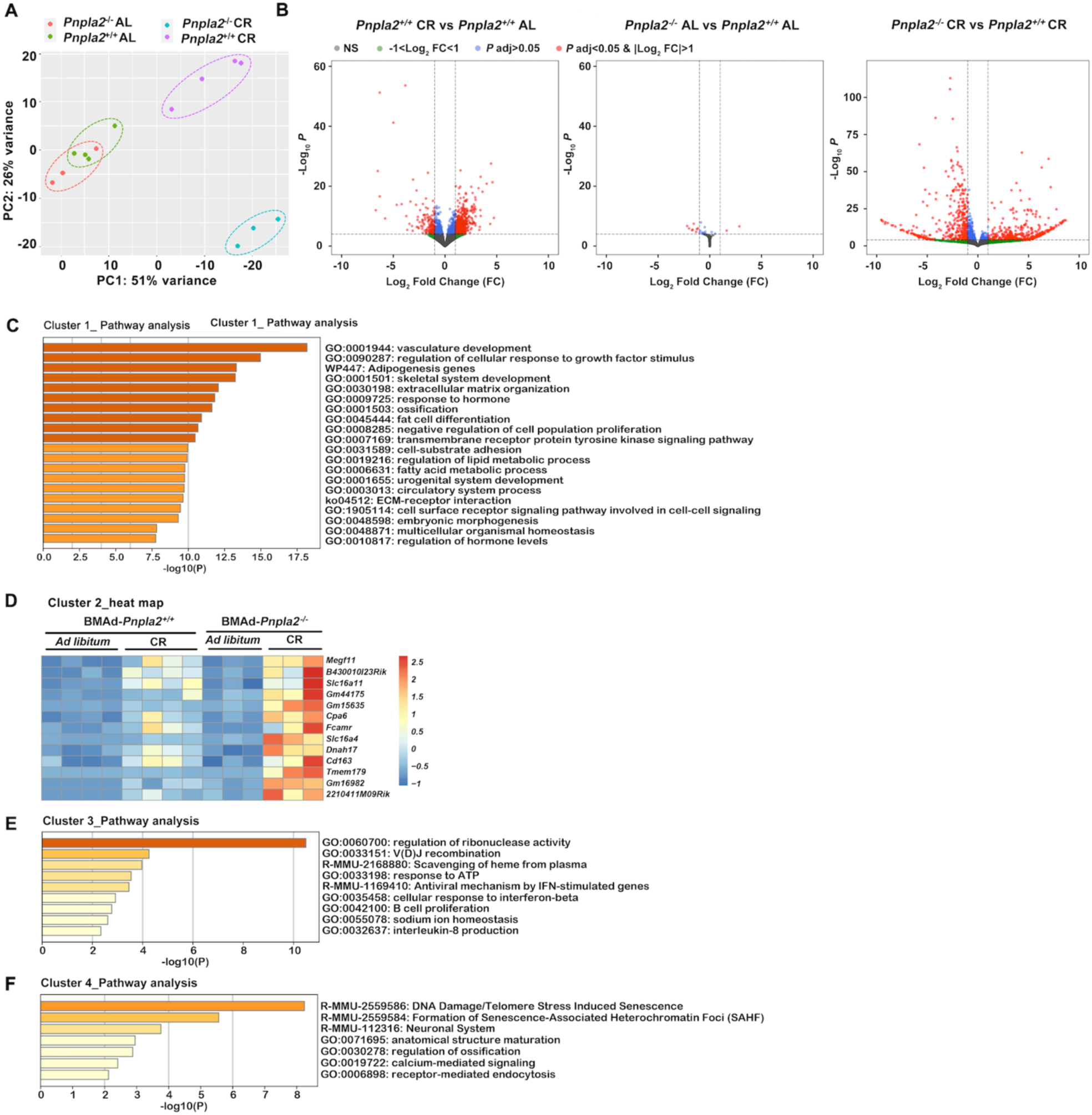
CR causes profound changes in BMAT transcriptome. 24 weeks old male mice underwent 30% CR for 6 weeks. Distal tibial cBMAT was flushed and cBMAT from two mice were pooled as one sample for RNA sample preparation. High-quality RNA samples were submitted for RNAseq analyses. A. Principal component analysis (PCA) plot shows the distinct transcriptional characters in CR groups with (purple dots) or without (aqua dots) *Pnpla2* in BMAds. B. Volcano plots show the differential genes with P adj<0.05 & |Log2 fold change|>1 in comparisons between BMAd-*Pnpla2*^+/+^ CR versus AL (left), BMAd-*Pnpla2*^-/-^ versus BMAd-*Pnpla2*^+/+^ at AL (middle) and BMAd-*Pnpla2*^-/-^ versus BMAd-*Pnpla2*^+/+^ at CR (right). C-F. Differential genes from comparison between BMAd-*Pnpla2*^+/+^ CR versus AL were grouped into 4 clusters according to the alteration patterns. Pathway analyses were performed on each cluster except cluster 2, which gene set was not enriched in any pathways.

**Figure 5 - figure supplement 2.**
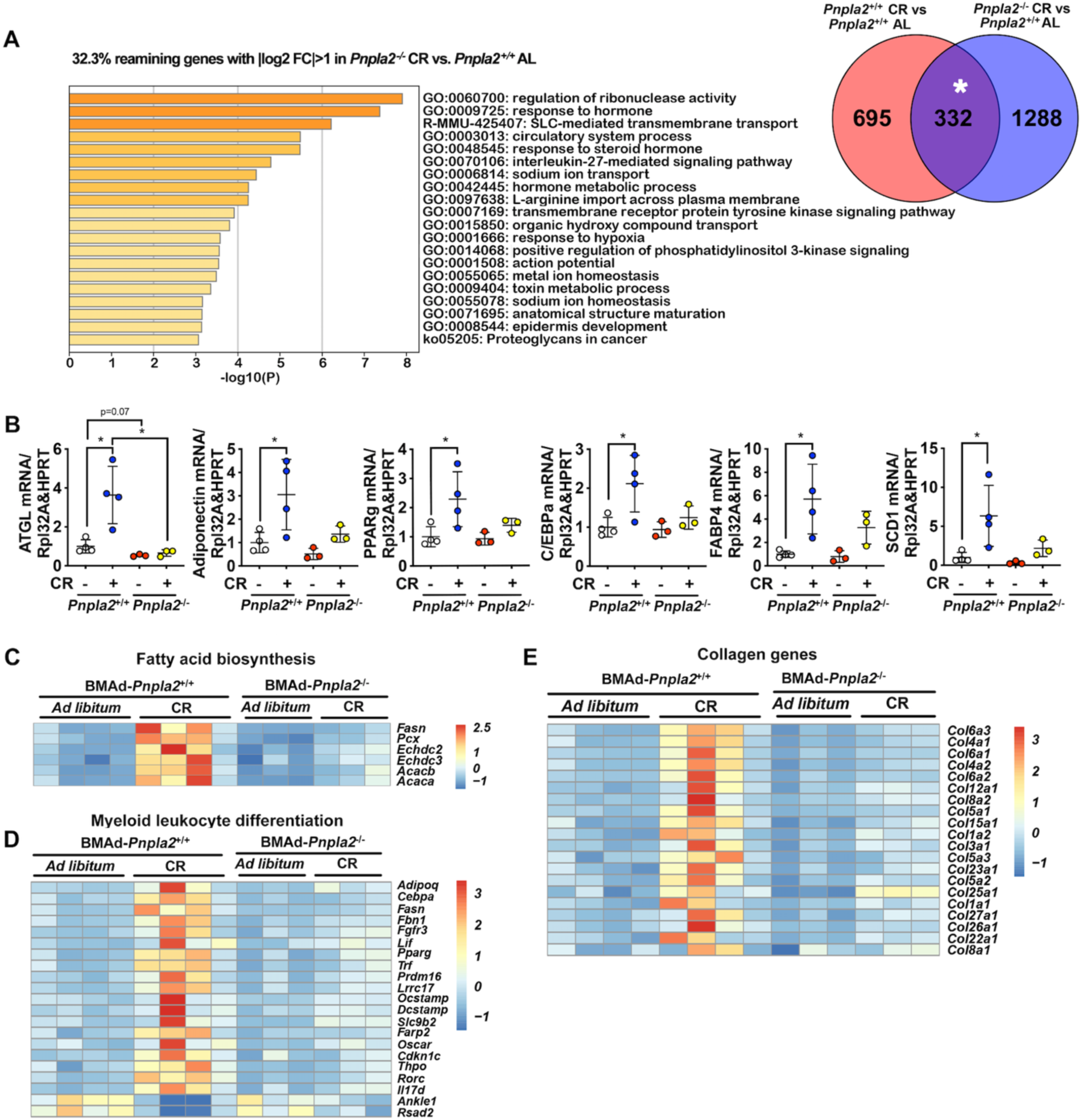
BMAd-*Pnpla2* deficiency causes extensive alterations to the bone marrow transcriptome only when coupled with CR. 24 weeks old male mice underwent 30% CR for 6 weeks. Distal tibial cBMAT was flushed and cBMAT from two mice were pooled as one sample for RNA sample preparation. High-quality RNA samples were submitted for RNAseq analyses. A. Pathway analysis of gene set that respond to CR independent of *Pnpla2* deficiency in BMAds, indicated by * area. B. qPCRs were performed to confirm the changes of adipogenesis genes in BMAT. Data are expressed as mean ± SD. * indicates *P* < 0.05 with one-way ANOVA with Tukey’s multiple comparisons test. C-D. Heat maps for genes related to fatty acid biosynthesis and myeloid leukocyte differentiation. E. Collagen genes that upregulated by CR in BMAd-*Pnpla2*^+/+^ mice.

