## Supplementary material for "Lipolysis of bone marrow adipocytes is required to fuel bone and the marrow niche during energy deficits": Table 1

**Table 1. BMAd-lipolysis deficiency has subtle effects on circulating mature blood cells.**

***Ad libitum*** --- Complete Blood Count (CBC)

| | | ***BMAd-ATGL*** $^{+/+}$ ***(n=5)*** | | ***BMAd-ATGL*** $^{-/-}$ ***(n=6)*** | | |
|---|---|---|---|---|---|---|
| *Parameter* | Units | Mean | Std | Mean | Std | P values |
| *WBC* | K/uL | 6.39 | 0.90 | 5.04 | 2.53 | 0.33 |
| *NE#* | K/uL | 2.58 | 1.25 | 2.03 | 0.60 | 0.41 |
| *LY#* | K/uL | 3.28 | 1.29 | 2.64 | 1.97 | 0.58 |
| *MO#* | K/uL | 0.43 | 0.21 | 0.30 | 0.12 | 0.26 |
| *EO#* | K/uL | 0.08 | 0.10 | 0.07 | 0.06 | 0.78 |
| *BA#* | K/uL | 0.012 | 0.01 | 0.01 | 0.01 | 0.77 |
| *NE%* | % | 40.52 | 18.02 | 46.88 | 15.88 | 0.59 |
| *LY%* | % | 51.51 | 18.25 | 45.39 | 16.75 | 0.61 |
| *MO%* | % | 6.54 | 2.27 | 6.28 | 1.98 | 0.86 |
| *EO%* | % | 1.25 | 1.42 | 1.31 | 0.74 | 0.94 |
| *BA%* | % | 0.18 | 0.15 | 0.14 | 0.13 | 0.67 |
| *RBC* | M/uL | 9.15 | 1.02 | 8.77 | 0.98 | 0.59 |
| *HB* | g/dL | 14.04 | 1.92 | 13.53 | 1.56 | 0.67 |
| *HCT* | % | 42.68 | 5.60 | 41.32 | 4.55 | 0.70 |
| *MCV* | fL | 46.54 | 1.91 | 47.17 | 1.93 | 0.64 |
| *MCH* | Pg | 15.32 | 0.82 | 15.47 | 0.91 | 0.81 |
| *MCHC* | g/dL | 32.88 | 1.03 | 32.75 | 1.10 | 0.86 |
| *RDW* | % | 19.02 | 0.90 | 18.63 | 0.92 | 0.54 |
| *PLT* | K/uL | 864.8 | 127.15 | 933 | 121.51 | 0.43 |
| *MPV* | fL | 5.18 | 0.63 | 4.7 | 0.51 | 0.24 |

**Caloric restriction** --- CBC

| | | ***BMAd-ATGL*** $^{+/+}$ ***(n=7)*** | | ***BMAd-ATGL*** $^{-/-}$ ***(n=7)*** | | |
|---|---|---|---|---|---|---|
| *Parameter* | Units | Mean | Std | Mean | Std | P values |
| *WBC* | K/uL | 3.98 | 0.93 | 3.56 | 1.02 | 0.43 |
| *NE#* | K/uL | 1.80 | 0.52 | 1.19 | 0.39 | **0.03*** |
| *LY#* | K/uL | 1.89 | 0.78 | 2.13 | 0.67 | 0.54 |
| *MO#* | K/uL | 0.15 | 0.08 | 0.18 | 0.05 | 0.44 |
| *EO#* | K/uL | 0.12 | 0.10 | 0.05 | 0.06 | 0.13 |
| *BA#* | K/uL | 0.03 | 0.03 | 0.01 | 0.02 | 0.31 |
| *NE%* | % | 45.88 | 13.18 | 33.46 | 6.12 | **0.04*** |
| *LY%* | % | 46.94 | 14.20 | 59.62 | 6.87 | 0.05 |
| *MO%* | % | 3.70 | 1.04 | 5.30 | 1.40 | **0.03*** |
| *EO%* | % | 2.86 | 2.04 | 1.25 | 1.57 | 0.12 |
| *BA%* | % | 0.62 | 0.64 | 0.36 | 0.54 | 0.43 |
| *RBC* | M/uL | 9.21 | 0.98 | 9.74 | 0.80 | 0.29 |
| *HB* | g/dL | 13.04 | 1.43 | 13.84 | 1.22 | 0.28 |
| *HCT* | % | 42.57 | 4.47 | 45.16 | 3.70 | 0.26 |
| *MCV* | fL | 46.24 | 1.82 | 46.37 | 0.92 | 0.87 |
| *MCH* | Pg | 14.16 | 0.60 | 14.21 | 0.52 | 0.85 |
| *MCHC* | g/dL | 30.63 | 0.66 | 30.66 | 0.79 | 0.94 |
| *RDW* | % | 18.56 | 0.95 | 18.09 | 0.41 | 0.25 |
| *PLT* | K/uL | 947.86 | 124.44 | 838.29 | 185.96 | 0.22 |
| *MPV* | fL | 5.13 | 0.52 | 4.89 | 0.37 | 0.33 |

Whole blood from male mice at 24 weeks of age fed *ad libitum* (top) or a 30% CR diet for 6 weeks (bottom) was submitted for complete blood counts (CBC). White- and red- blood cell related parameters are shown.
