## Supplementary material for "Lipolysis of bone marrow adipocytes is required to fuel bone and the marrow niche during energy deficits": Table 2

**Table 2. BMAd-lipolysis deficiency causes mild changes in bone marrow hematopoietic cells.**

***Ad libitum*** --- Flowcytometry

| | | *BMAd-ATGL$^{+/+}$ (n=5)* | | *BMAd-ATGL$^{-/-}$ (n=6)* | | | | *BMAd-ATGL$^{+/+}$ (n=5)* | | *BMAd-ATGL$^{-/-}$ (n=6)* | | |
|---|---|---|---|---|---|---|---|---|---|---|---|---|
| *Counts* | Units | Mean | Std | Mean | Std | P values | Percentage | Mean | Std | Mean | Std | P values |
| *BMNC* | 10^6/ml | 21.89 | 1.78 | 19.33 | 3.27 | 0.31 | \ | \ | \ | \ | \ | \ |
| *NE#* | 10^6/ml | 13.93 | 1.39 | 11.63 | 1.73 | 0.17 | NE% | 63.62 | 3.60 | 60.43 | 3.66 | 0.18 |
| *MO#* | 10^5/ml | 0.74 | 0.23 | 0.69 | 0.32 | 0.90 | MO% | 0.34 | 0.09 | 0.35 | 0.16 | 0.83 |
| *B cell* | 10^5/ml | 5.96 | 4.46 | 8.55 | 7.56 | 0.98 | B cell% | 2.86 | 2.40 | 4.11 | 3.53 | 0.52 |
| *T cell* | 10^5/ml | 8.46 | 3.82 | 5.06 | 2.28 | 0.33 | T cell% | 3.8 | 1.53 | 2.68 | 1.35 | 0.23 |
| **HSPCs** | | | | | | | | | | | | |
| *HSC* | 10^3/ml | 1.88 | 1.35 | 1.58 | 0.96 | 0.67 | HSC% | 0.008 | 0.005 | 0.009 | 0.006 | 0.88 |
| *MPP* | 10^3/ml | 2.84 | 1.80 | 1.96 | 1.16 | 0.35 | MPP% | 0.013 | 0.008 | 0.010 | 0.007 | 0.59 |
| *GMP* | 10^3/ml | 5.36 | 2.44 | 1.85 | 1.48 | **0.02*** | GMP% | 0.024 | 0.011 | 0.010 | 0.008 | **0.03*** |
| *PreGM* | 10^3/ml | 8.59 | 6.78 | 7.38 | 5.06 | 0.74 | PreGM% | 0.039 | 0.031 | 0.039 | 0.026 | 0.99 |
| *PreMegE* | 10^3/ml | 2.38 | 2.31 | 2.00 | 1.34 | 0.74 | PreMegE% | 0.011 | 0.010 | 0.011 | 0.008 | 0.98 |
| *PreCFUe* | 10^3/ml | 0.35 | 0.09 | 0.58 | 0.57 | 0.41 | PreCFUe% | 0.002 | 0.0004 | 0.003 | 0.003 | 0.32 |
| *HPC 1* | 10^3/ml | 5.16 | 3.68 | 2.83 | 1.77 | 0.20 | HPC 1% | 0.025 | 0.019 | 0.014 | 0.007 | 0.23 |
| *HPC 2* | 10^3/ml | 1.01 | 0.64 | 0.39 | 0.24 | 0.05 | HPC 2% | 0.005 | 0.003 | 0.002 | 0.001 | 0.08 |

**Caloric restriction** --- Flowcytometry

| | | *BMAd-ATGL$^{+/+}$ (n=6)* | | *BMAd-ATGL$^{-/-}$ (n=7)* | | | | *BMAd-ATGL$^{+/+}$ (n=6)* | | *BMAd-ATGL$^{-/-}$ (n=7)* | | |
|---|---|---|---|---|---|---|---|---|---|---|---|---|
| *Counts* | Units | Mean | Std | Mean | Std | P values | Percentage | Mean | Std | Mean | Std | P values |
| *BMNC* | 10^6/ml | 20.79 | 6.00 | 16.64 | 2.44 | 0.12 | | \ | \ | \ | \ | \ |
| *NE#* | 10^6/ml | 9.40 | 3.25 | 5.83 | 1.44 | **0.02*** | NE% | 45 | 4.49 | 34.74 | 5.58 | **0.004*** |
| *MO#* | 10^5/ml | 1.43 | 0.44 | 1.12 | 0.24 | 0.14 | MO% | 6.89 | 0.99 | 6.71 | 0.88 | 0.73 |
| *B cell* | 10^5/ml | 41.96 | 21.14 | 52.68 | 10.73 | 0.26 | B cell% | 19.83 | 8.01 | 31.9 | 6.84 | **0.01*** |
| *T cell* | 10^5/ml | 10.09 | 6.64 | 7.10 | 1.578 | 0.27 | T cell% | 5.11 | 3.49 | 4.36 | 1.23 | 0.60 |
| **HSPCs** | | | | | | | | | | | | |
| *HSC* | 10^3/ml | 1.166 | 0.67 | 0.77 | 0.21 | 0.17 | HSC% | 0.057 | 0.032 | 0.05 | 0.015 | 0.50 |
| *MPP* | 10^3/ml | 0.93 | 0.31 | 0.79 | 0.20 | 0.37 | MPP% | 0.045 | 0.010 | 0.05 | 0.009 | 0.63 |
| *GMP* | 10^3/ml | 6.91 | 3.40 | 4.47 | 1.11 | 0.10 | GMP% | 0.33 | 0.114 | 0.27 | 0.046 | 0.21 |
| *PreGM* | 10^3/ml | 16.57 | 4.85 | 14.72 | 3.48 | 0.44 | PreGM% | 0.812 | 0.170 | 0.88 | 0.174 | 0.47 |
| *PreMegE* | 10^3/ml | 4.95 | 3.48 | 3.94 | 1.21 | 0.49 | PreMegE% | 0.248 | 0.178 | 0.24 | 0.077 | 0.91 |
| *PreCFUe* | 10^3/ml | 5.28 | 2.81 | 4.28 | 1.04 | 0.40 | PreCFUe% | 0.247 | 0.066 | 0.27 | 0.094 | 0.68 |
| *HPC 1* | 10^3/ml | 6.76 | 4.32 | 5.25 | 1.66 | 0.41 | HPC 1% | 0.33 | 0.202 | 0.33 | 0.136 | 0.98 |
| *HPC 2* | 10^3/ml | 16.08 | 12.78 | 10.94 | 2.90 | 0.32 | HPC 2% | 0.8 | 0.652 | 0.68 | 0.255 | 0.67 |

Femoral bone marrow cells from male mice at 24 weeks of age fed *ad libitum* (top) or a 30% CR diet for 6 weeks (bottom) were collected and stained with antibodies for flow cytometry analyses. Mature blood cells and hematopoietic stem/progenitor cells (HSPCs) were counted.
