## Supplementary material for "Lipolysis of bone marrow adipocytes is required to fuel bone and the marrow niche during energy deficits": Table 3

### Table 3. PCR primer list

| Name | Sequence |
|---|---|
| *Genotype PCR* | |
| *Osterix* p1 | CCGCCCCGATCTTCCACT |
| *Osterix* p2 | GTTGCCGGTCCTGTTCACTCTC |
| *Osterix* p3 | TGCTCCCGGCCAGGTTACTA |
| FAC p1 | CACTCTAAATGAACACGTGCTTTCG |
| FAC p2 | ACCGGCATCAACGTTTTCTTTT |
| FAC p3 | CTGGATAGTGAAACAGGGGCA |
| FAC-WT Forward | AGCCCATACACCAGGAGAATCA |
| FAC-WT Reverse | TGTGAAGCCCCCATACCAA |
| FAC-Mut Forward | AGCCCATACACCAGGAGAATCA |
| FAC-Mut Reverse | TTGGCGAGAGGGGAAAGACC |
| *Regular PCR* | |
| *Pnpla2* Forward | CCAACGCCACTCACATCTACG |
| *Pnpla2* Reverse | ACCCCGGGGCTCCTCTTA |
| *qPCR* | |
| FAC Original band- Forward | TCGCTAGCTCAATCGCCATCTT |
| FAC Original band-Reverse | GCCACCAGCCAGCTATCAACTC |
| FAC Flipped band- Forward | TGAAGGATGCCCAGAAGGTA |
| FAC Flipped band-Reverse | CGGCAAACGGACAGAAGC |
| *Pnpla2* Forward | CGGCTTCCTCGGGGTCTAC |
| *Pnpla2* Reverse | CGCGCTCATGGCAATCAG |
| *Adipoq* Forward | CATTCCGGGACTCTACTACTTCT |
| *Adipoq* Reverse | GAGGCCTGGTCCACATTCTT |
| *Ppar$\gamma$* Forward | GCCATTGAGTGCCGAGTCTGT |
| *Ppar$\gamma$* Reverse | GCATCCGCCCAAACCTGA |
| *Cebp$\alpha$* Forward | TGGACAAGAACAGCAACGAG |
| *Cebp$\alpha$* Reverse | TCACTGGTCAACTCCAGCAC |
| *Fabp4* Forward | ATGAAATCACCGCAGACGACA |
| *Fabp4* Reverse | CACGCCTTTCATAACACATTCC |
| *Scd1* Forward | CGTGGGTTGGCTGCTTGTG |
| *Scd1* Reverse | CAGGAGGCCGGGCTTGTAGT |
| *Rpl32a* Forward | GAGCAACAAGAAAACCAAGCA |
| *Rpl32a* Reverse | TGCACACAAGCCATCTACTCA |
| *Hprt* Forward | TCATTATGCCGAGGATTTGGA |
| *Hprt* Reverse | GCACACAGAGGGCCACAAT |
| *Tbp* Forward | ACCTTATGCTCAGGGCTTGG |
| *Tbp* Reverse | GCCGTAAGGCATCATTGGAC |
